# Mitochondrial Matrix Protease ClpP Agonists inhibit Cancer Stem Cell Function in Breast Cancer Cells by Disrupting Mitochondrial Homeostasis

**DOI:** 10.1101/2022.03.18.484947

**Authors:** Yoshimi E. Greer, Lidia Hernandez, Emily M. J. Fennell, Donna Voeller, Raj Chari, Sam Gilbert, Binwu Tang, Markus Hafner, Christina M. Annunziata, Edwin Iwanowicz, Lee M. Graves, Stanley Lipkowitz

**Affiliations:** Women’s Malignancies Branch, NCI, NIH, Bethesda, MD; Department of Pharmacology, University of North Carolina School of Medicine, Chapel Hill, NC; Genome Modification Core, Frederick National Laboratory for Cancer Research, NCI, NIH, Frederick, MD; Laboratory of Cancer Biology and Genetics, NCI, NIH; RNA Molecular Biology Group, Laboratory of Muscle Stem Cells and Gene Regulation, NIAMS, NIH, Bethesda, MD; Madera Therapeutics, LLC, Cary, NC

## Abstract

Mitochondria are multifaceted organelles which are important for bioenergetics, biosynthesis and signaling in metazoans. Mitochondrial functions are frequently altered in cancer to promote both the energy and the necessary metabolic intermediates for biosynthesis required for tumor growth. Cancer stem cells (CSCs) contribute to chemotherapy resistance, relapse, and metastasis. Recent studies have shown that while non-stem, bulk cancer cells utilize glycolysis, breast CSCs are more dependent on oxidative phosphorylation (OxPhos) and therefore targeting mitochondria may inhibit CSC function. We previously reported that small molecule ONC201, which is an agonist for the mitochondrial caseinolytic protease (ClpP), induces mitochondrial dysfunction in breast cancer cells. In this study, we report that ClpP agonists inhibit breast cancer CSC function *in vitro* and *in vivo*. Mechanistically, we found that OxPhos inhibition downregulates multiple pathways required for CSC function, such as the mevalonate pathway, YAP, Myc, and the HIF pathway. ClpP agonists showed significantly greater inhibitory effect on CSC functions compared with other mitochondria-targeting drugs. Further studies showed that ClpP agonists deplete NAD(P)+ and NAD(P)H and induce redox imbalance, and dysregulate one-carbon metabolism and proline biosynthesis. Downregulation of these pathways by ClpP agonists further contribute to the inhibition of CSC function. In conclusion, ClpP agonists inhibit breast CSC functions by disrupting mitochondrial homeostasis in breast cancer cells and inhibiting multiple pathways critical to CSC function.

## Introduction

Mitochondria regulate multiple cell functions including energy metabolism, cell death and survival, and signaling pathways(1). Mitochondria are broadly implicated in cancer biology, and deregulation of cellular energetics has been recognized as a hallmark of cancer(2).

Previously, we reported that the small molecule compound ONC201 inhibits cell viability of breast cancer cells by targeting mitochondria(3). ONC201 inhibited OxPhos, depleted cellular ATP, and induced a stress response. Subsequent studies demonstrated that ONC201 binds and activates mitochondrial caseinolytic protease ClpP(4,5), a serine protease located in the mitochondrial matrix. ClpP maintains mitochondrial protein homeostasis by degrading misfolded or damaged proteins(6). The known substrates of ClpP include proteins essential for the electron transport chain, the tricarboxylic acid cycle, mitochondrial gene transcription and translation(7). Thus, dysregulation of mitochondrial homeostasis by ClpP agonists is considered a novel strategy for cancer treatment(6).

The existence of CSCs is a major obstacle for cancer therapy because CSCs contribute to drug resistance, relapse, and metastasis(8). Mitochondrial function and energy metabolism are important factors to sustain the stemness of CSCs(9). Studies have shown that breast CSCs are more dependent on OxPhos, while differentiated, proliferative progeny display a glycolytic phenotype(10,11). Therefore, eliminating CSCs via targeting mitochondrial function has the potential to improve long-term outcomes in breast cancer(9,12,13).

In this study, we examined the effect of ClpP agonists on CSC function in breast cancer. ClpP agonists inhibited tumor initiating ability *in vitro* and *in vivo*. Mechanistically, ClpP agonists dysregulated multiple signaling pathways involved with CSC functions, including the mevalonate, YAP, Myc, and HIF pathways. While other mitochondrial-targeted drugs also inhibited these pathways, ClpP agonists showed significantly greater impact on CSC functions compared with other drugs. ClpP agonists also depleted coenzymes NAD(P)+ and NAD(P)H and induced redox imbalance, essential factors to maintain CSCs. Further, ClpP agonist uniquely downregulated multiple mitochondrial enzymes involved with folate-mediated one-carbon metabolism (FOCM) and proline biosynthesis. Finally, we found that proline biosynthesis is required for breast CSC functions. In conclusion, ClpP agonists inhibit breast CSCs by not only targeting OxPhos but disrupting multiple metabolic pathways essential for CSC function.

## Methods and Materials

(See Supplementary Table S1 for detailed reagent and equipment information)

### Reagents

ONC201 was provided by Chimerix, Inc., and TR-compounds were provided by Madera Therapeutics, LLC. For other reagents, equipment and software see Table S1.

### Cell Culture

Human breast cancer cell lines MDA-MB-231 (MB231), MCF7, MDA-MB-453 (MB453), SKBR3 cells were obtained from ATCC (Manassas, VA), and maintained in RPMI1640 supplemented with 10 % fetal bovine serum (FBS), 100 units/ml of penicillin, 100 *μ*g/ml of streptomycin (P/S). SUM159 CLPP WT and KO cells were gifts from Dr. Lee Graves, UNC, and maintained with DMEM/F12 supplemented with 5% FBS, 100 *μ*g/mL of P/S, 5 µg/mL insulin and 1 µg/ml hydrocortisone. The MB231* rho0 (mtDNA depleted) cell line was maintained as previously reported(3). MB231 LM2 SORE6-mCherry-CD19+ and MB231 LM2 mCMV-mCherry (Ctl.) were provided by Dr. Lalage Wakefield, NCI and maintained with DMEM supplemented with 10 % FBS(14). All cells were maintained at 37 °C, 5 % CO_2_ incubator.

### Acridine orange / Propidium iodide (AO/PI) cell viability assay

Live/dead and total cell numbers were counted with AOPI and Cellometer K2 (See Table S1).

### Mammosphere formation assays

Mammosphere formation was measured as previously described with slight modification(15). Briefly, DMEM/F12 supplemented with SingleQuots was used as basal assay media. B-27 supplement and bFGF were freshly added to the basal media for non-adherent cell culture condition. To examine the effect of drugs on sphere formation, three different protocols were used. In the first protocol, adherent cells pretreated with drugs for 48 h, were then trypsinized, rinsed with PBS, and single cell suspensions were prepared with the basal media. From these suspensions 1,000 viable cells were seeded per well of low-attachment chamber with mammosphere assay media in triplicate. After 10-14 days, the numbers of mammosphere (diameter more than 50 μm) were counted using a standard microscope with 4x or 10x magnification. For the second and third protocols, cells were directly seeded on to low-attachment chambers at a density of 1,000 cells/well in mammosphere assay media, and the drugs were directly added to the media on the next day (one application) or multiple times (every 2-3 days).

### Energy metabolism assays using bioluminescence

CellTiter-Glo® 2.0, RealTime-Glo™ MT Cell Viability, NAD(P)/NAD(P)H-Glo™, ROS-Glo^™^ H_2_O_2_, GSH/GSSH-Glo™ assays were all performed according to the manufacturer’s protocol. Luminescence was measured by a SpectraMax^R^ iD3 microplate reader. All measurements were performed in triplicate and each experiment was carried out at least 3 times.

### Colorimetric Proline assay and Glucose 6 Phosphate Dehydrogenase (G6PDH) activity assay

The proline level in cells was analyzed using a General Proline Assay Kit. A G6PDH assay kit was used to measure G6PDH activity. SpectraMax^R^ iD3 microplate reader was used for both assays.

### Generation of MB231 CLPP WT and KO cell lines using CRISPR/Cas9 system

Candidate single guide RNAs (sgRNAs) targeting the *CLPP* gene were identified using the sgRNA Scorer 2.0 design tool(16). Eight candidate sgRNAs were tested for cutting activity in HEK293T cells, and sgRNA1 and sgRNA2 were used for experiments in breast cancer cells. Oligonucleotides encoding for these guides, along with a non-targeting control guide RNA, were obtained from IDT Technologies and subsequently annealed and ligated into the Lenti-CRISPR-V2 backbone using T4 ligation. Ligated products were then transformed into Stbl4 competent cells. LentiCRISPR v2 was a gift from Dr. Feng Zhang, MIT, Broad Institute (17). The plasmids grown in bacteria were purified with EndoFree Plasmid Maxi Kit. Lipofectamine 3000 was used for DNA transfection. Forty-eight hours after transfection, puromycin was added to cell culture (1 μg/ml) for one week. Cells were trypsinized and re-seeded on to 96 well plates at density of one cell/well to establish single clonal lines. Clones grown were assessed for *CLPP* KO status by PCR from the genomic DNA and subsequent deep amplicon Illumina sequencing encompassing the target sites, as described(16). The status of CLPP KO in MB231 and MCF7 cell lines were analyzed using a custom computational pipeline to determine editing rate in each clone (Fig.S1A, Table S1). Loss of protein expression was further confirmed by Western blotting.

### siRNA transfection, mtDNA copy number, reverse transcriptase-quantitative PCR, Western blotting

All were performed as reported previously(3).

### Cell fractionation

All were performed as described elsewhere(18).

### ALDEFLUOR assay

ALDEFLUOR™ Kit and ALDEFLUOR™ DEAB Reagent (negative control) were used according to the manufacturer’s protocol. After incubation for indicated drug treatment, cells were trypsinized and cell suspensions (1×10^6^ cells/ml) were prepared for each condition. Once ALDEFLUOR™ reagent and LIVE/DEAD Aqua were added, cells were incubated for 45 minutes at 37 °C, washed twice, resuspended with 0.5 ml the assay buffer, and filtered using Cell Strainer prior to analysis with a BD FACS Verse Flow Cytometer. Data was analyzed with FlowJo software.

### Sox2/Oct4 responsive element (SORE) promoter-driven stem cell reporter assay (SORE6 reporter assay

MB231-LM2-mCMV-mCherry (control) and MB231-LM2-SORE6-mCherry cells were plated at 1×10^5^ cells/well in 6-well plates. After indicated times of drug treatment, cells were trypsinized, collected, washed in PBS, and diluted to 1×10^6^ cells/mL for staining with LIVE/DEAD® Fixable Blue Dead Cell Stain. Cells were then centrifuged for 5 minutes at 100x*g* in 4 °C, then washed with PBS, and resuspended in FACS buffer (4 % FBS/PBS with 0.5 mM EDTA). Cells were passed through a cell strainer for FACS analysis with BD Fortessa. MB231-LM2-mCMV-mCherry cells were defined as SORE6 positive if the fluorescence in the mCherry exceeded that of 99.9 % of the MB231-LM2-mCMV control line as previously described(14).

### HOPflash, HRE-Luc reporter assays

Cells were seeded in 96 well white plates one day prior to transfection. Reporter genes HOPflash or HRE-Luc were transfected with the internal control NanoLuc (pNL1.1.TK) using Lipofectamine 3000. After 24 h, cells were treated with indicated drugs for 48 h. Luciferase was measured using the Nano-Glo® Dual-Luciferase® Reporter Assay System and SpectraMax^R^ iD3 microplate reader.

### Seahorse XF Real-Time ATP Rate Assay

Oxygen Consumption Rate (OCR), Extra Cellular Acidification Rate (ECAR), and OxPhos-ATP or Glycolysis-ATP were measured with a XFe24 Extracellular Flux Analyzer with FluxPaks Mini and using a XF Real-Time ATP Rate Assay Kit. Cells were seeded on a XFe24 cell culture microplate (6×10^4^ cells/well) with growth medium, and on the following day, cells were treated with the indicated drugs and incubated for 24 h. On the day of the assay, the medium was replaced with XF assay buffer (DMEM pH 7.4, 10 mM glucose, 1 mM pyruvate, 2 mM glutamine). An ATP rate assay was performed per the manufacturer’s instructions. After the assay, the cells were fixed with 3.7% formaldehyde/PBS for 15 min, washed with PBS twice and stained with Hoechst (1 μg/ml in PBS), and cell numbers were counted with Cytation 1 for normalization. Analysis of the ATP rate assay was performed with manufacturer’s software. The ATP rate index was calculated as the ratio of OxPhos-ATP/Glycolysis-ATP.

### XF assays with mammospheres

To enable attachment of cells from mammospheres, XFe24 cell culture microplates were pre-coated with poly-L-lysine (100 μg/ml), 50 μl/well, then incubated at 37 °C for 2 h. The coated wells were rinsed with water 3 times, stored at 4 °C and used within one week. On the day of assay, pre-coated microplates were rinsed twice with PBS, and once with XF assay buffer. Mammospheres grown in mammosphere culture media were collected and single cell suspensions were prepared by pipetting. Cell numbers were counted with AOPI assays, and 6×10^4^ viable cells were seeded with 100 μl assay buffer/well of the pre-coated microplates. The assay plates were centrifuged at 200xg for 1 minute, transferred to 37 °C CO_2_ incubator for 30 minutes to ensure cell attachment, 400 μl of XF assay buffer/well was overlaid (total 500 μl/well), and immediately used for assays.

### Tumorigenicity assay in vivo

The effect of ClpP targeting on tumor initiation capacity *in vivo* was evaluated in two animal experiments. In both experiments, 6-8 weeks old athymic nude female mice were used. In the first experiment, MB231 cells were first treated with ONC201 (5 μM) or DMSO (Ctl.) for 48 h *in vitro*. Cells were trypsinized, collected, and viable cell number was determined with AOPI. Cell suspensions were prepared with 2 different cell densities (5×10^5^, or 5×10^6^ viable cells/mouse), suspended in PBS (50μL/mouse), and injected into mammary fat pad (MFP)(10 mice per arm). The animals were not treated with drug. Tumor formation, tumor size, and body weight were followed for up to 43 days. In the second experiment, MB231 CLPP WT or KO cells were treated with either TR-57 (50 nM) or DMSO (Ctl.) for 48 h. Cells were collected and counted as above, 5×10^5^ or 5×10^6^ cells in PBS were injected into MFP, 10 mice per arm in the absence of drug. Tumor formation was monitored for up to 48 days. CSC frequency was calculated with Extreme Limiting Dilution Analysis (ELDA) software. The tumor volume was calculated using formula: tumor volume (mm^3^) = length x (width)^2^ x 0.5. Animal maintenance and experiments were performed in accordance with the animal care guidelines of NIH, Bethesda, MD, USA. All animal experiments were approved by the Animal Research Advisory Committee of NCI, NIH, Bethesda, MD.

### Isolation of xenografted cells from tumors by depletion of mouse cells

Tumors grown in mice in the first tumorigenicity experiment were collected and human cells were isolated by depleting mouse cells, as described in the manufacturer’s protocol (see Table S1).

### Network and pathway analysis

Bioinformatics analysis of RNAseq data of MB231 treated with ONC201 was performed as previously reported(3). The source of Gene Set Enrichment Analysis (GSEA), Ingenuity Pathway Analysis (IPA), MetaCore are shown in Table S1.

### Statistical analysis

The significance of differences in data was determined with Student’s *t*-test, or one-way ANOVA, or 2-way ANOVA as indicated in figure legends. The differences were considered significant when *p* value was less than 0.05.

## Results

### ClpP agonists inhibit cell viability and OxPhos in breast cancer cells in a CLPP-dependent manner

We tested multiple ClpP agonists (ONC201, TR-57, TR-65) in breast cancer cell lines (Fig.1A). All ClpP agonists decreased cell viability in multiple breast cancer cells, including MB231, MB453, MCF7 (Fig.1B). TR compounds were ∼60 to 270-fold more potent (Fig.1C) compared with ONC201 in all cell lines tested. Consistent with our previous observation that ONC201 depletes Tfam proteins and mtDNA(3), all ClpP agonists depleted Tfam (Fig.1D) and depleted mtDNA (MB231: Fig.1E, MCF7: Fig.S1B).

**Fig.1.**
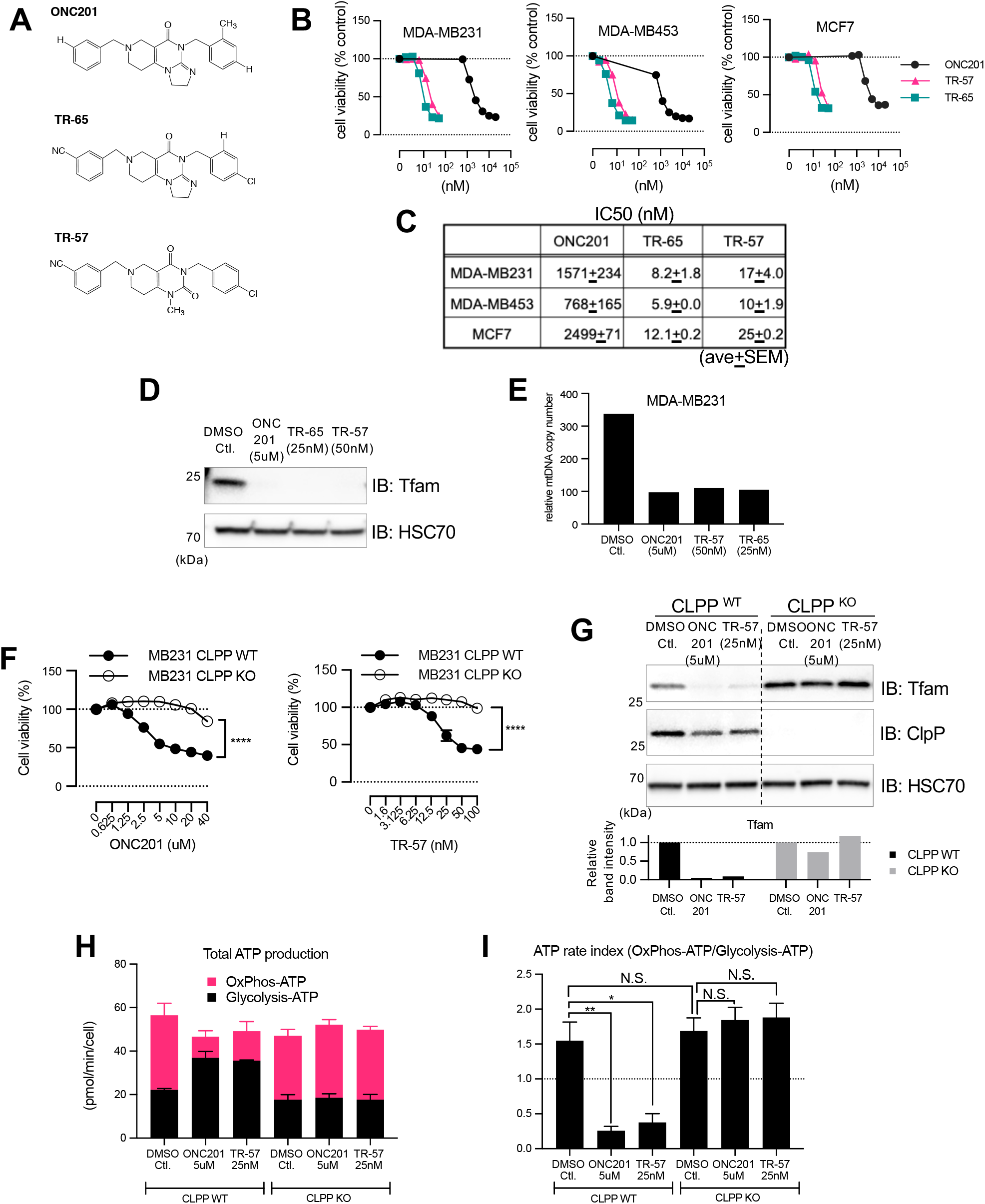
ClpP agonists inhibit cell viability and OxPhos in breast cancer cells in a CLPP-dependent manner. **A**. Chemical structures of ClpP agonists. **B**. CellTiter-Glo 2.0 assay performed after 72h treatment. Data shown as Ave+/-SEM of 3 independent experiments. **C**. IC50 (nM) of ClpP agonists in 3 breast cancer cell lines. **D**. Western blot of Tfam levels in control or ONC201 and TR-treated MB231 cells after 48h treatment. **E**. mtDNA copy number analyzed with quantitative PCR (48h post-treatment). **F**. CellTiter-Glo 2.0 assay in MB231 CLPP WT and CLPP KO cells. Data shown as Average+/-SEM of 3 independent experiments. \*\*\*\**p*<0.0001 with 2-way ANOVA. **G**. Western blot showing the effect of ONC201 and TR-57 (48h treatment) on Tfam in MB231 CLPP WT and KO cell lines. **H**. Seahorse XF analyzer ATP rate assay of MB231 CLPP WT and KO cell lines treated with DMSO Ctl, ONC201, or TR-57 for 24h. Data shown as ave+/-SEM of 3 independent experiments. No significant difference in total ATP production was detected among all groups (*t*-test). **I**. ATP rate index obtained with ATP rate assay shown in Fig.1I. Data shown as ave+/-SEM of 3 independent experiments. \*\**p*<0.01, \**p*<0.05, Student’s *t*-test.

Next, we tested target specificity of ONC201 and TR compounds using MB231 CLPP KO cells (Fig. S1A). Both ONC201 and TR-57 inhibited cell viability of CLPP WT cells, but not in CLPP KO cells (Fig.1F), confirming that cytotoxic effects of these drugs are dependent on CLPP. ClpP agonists directly impair OxPhos ATP production(3-5) measured by the CellTiter-Glo 2.0 assay. Therefore, we repeated these experiments using the ATP-independent RealTime-Glo MT assay, and similarly observed that ONC201 and TR-57 inhibited cell viability in a CLPP-dependent manner (Fig.S1C). The CLPP-dependent cytotoxicity effects of ONC201 and TR-57 were also confirmed in MCF7 and SUM159 cells (Fig.S1D and E). Similarly, both ONC201 and TR-57 depleted Tfam in CLPP WT cells but not in CLPP KO cells (MB231, Fig.1G, MCF7: Fig.S1F, SUM159: Fig.S1G). Moreover, TR-57 decreased mtDNA copy number in MB231 CLPP WT cells, but not in CLPP KO cells (Fig.S1H).

Both ONC201 and TR-57 impaired OxPhos-ATP production resulting in decreased ATP rate index (e.g., the ratio of OxPhos-ATP/Glycolysis-ATP) in CLPP WT but not KO cells in MB231 and SUM159 cells (Fig.1H&I and Fig.S1I&J, respectively). An increase in ATP produced by glycolysis was seen in MB231 but not in SUM159. Together, these data demonstrate that the effects of ONC201 and TR-57 are CLPP-dependent.

### Mitochondria are critical for mammosphere formation

We previously established mtDNA-depleted MB231 cell lines (MB231*rho0)(3). MB231*rho0 cells had no detectable mtDNA (Fig.S2A), and the OCR was significantly downregulated (Fig.S2B). In mammosphere formation assays, MB231*rho0 cells formed significantly fewer mammospheres compared to the parental MB231 cells (Fig.2A), suggesting that functional mitochondria are required for CSC function. Next, we compared mtDNA copy number and bioenergetic status between adherent cells and mammospheres. The mtDNA copy number was significantly increased in mammospheres compared with adherent cells (Fig.2B for MB231, Fig.S2C for MCF7). Total ATP production was not different between adherent cells and mammospheres, however, mammospheres had a higher fraction of OxPhos-ATP compared with adherent cells, supporting the hypothesis that breast CSCs are more dependent on OxPhos (MB231: Fig.2C&D, MCF7: Fig.S2D&E). Transcript levels of several CSC markers (e.g., CD44, Myc, EpCAM, ZEB) were increased in mammospheres compared to adherent cells (MB231: Fig.2E, MCF7: Fig.S2F). These data suggest that CSCs are dependent on mitochondria.

**Fig.2.**
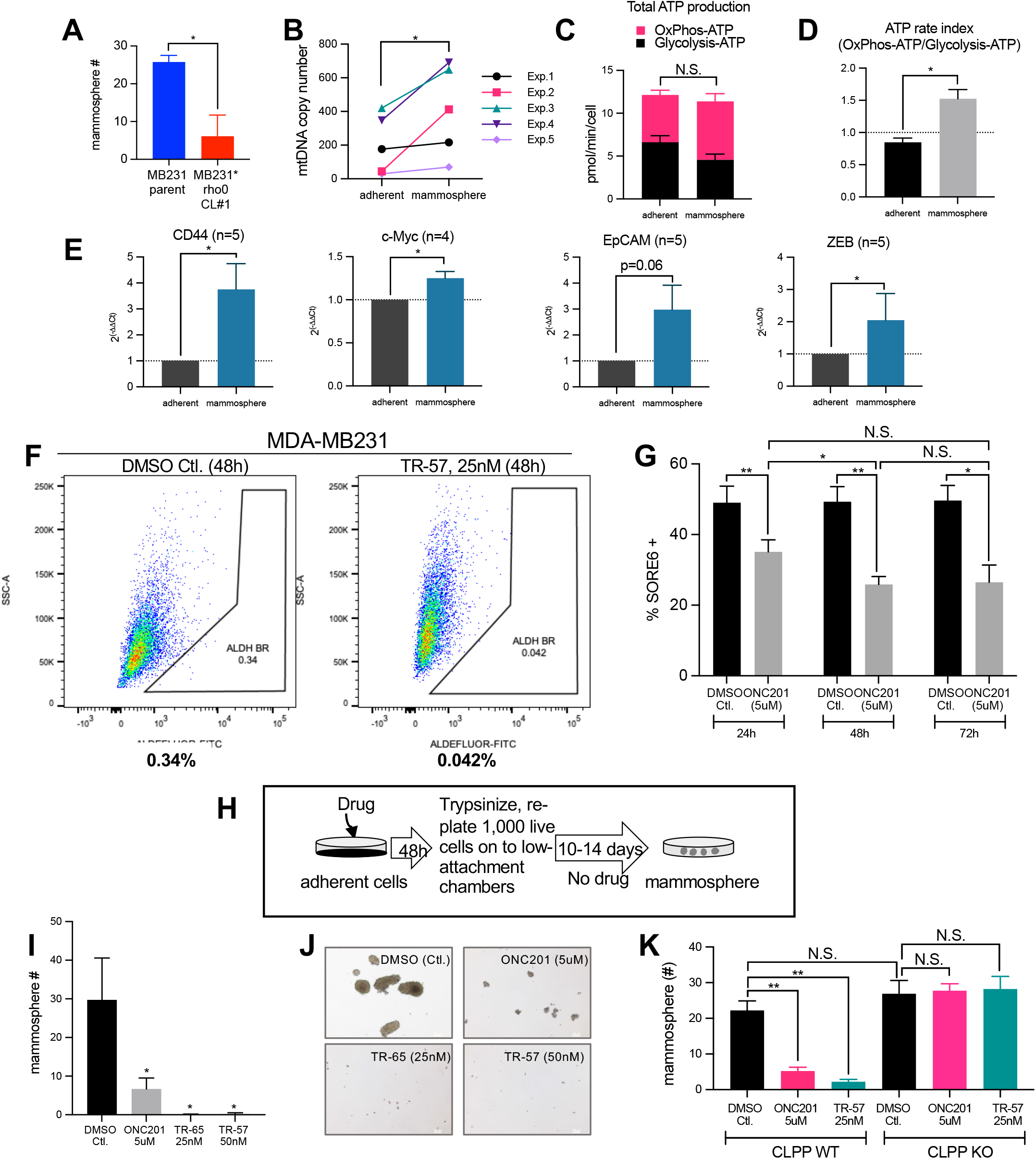
Mitochondria is critical for mammosphere formation and ClpP agonists inhibit CSC function *in vitro*. **A**. Mammosphere formation assays of MB231 parental cells and MB231* rho0 cells. Data shown as ave+/-SEM of 3 independent experiments. \**p*<0.05, Student’s *t*-test. **B**. mtDNA copy numbers between MB231 adherent cells and mammosphere. 5 independent experiments, \**p*<0.05, Paired *t*-test. **C**&**D**. Seahorse XF analyzer ATP rate assays of MB231 adherent cells and cells from mammospheres. Data shown as ave+/-SEM, summary of 3 independent experiments. \**p*<0.05, Student’s *t*-test. **E**. Stem cell markers mRNA measured using qPCR in MB231 adherent cells and mammospheres. Data shown as ave +/-SEM of multiple experiments. \**p*<0.05, Student’s t-test. **F**. Representative data of ALDEFLUOR assays of MB231 cells treated with DMSO or TR-57 (25nM) for 48h. Numbers shown below the panels indicate ALDH Bright considered as CSC population. **G**. MB231 SORE6 mCherry reporter assays. Cells were treated with ONC201 for indicated times. Data shown as ave+/-SEM, summary of 3 independent experiments. **H**. Diagram illustrating the procedure of mammosphere formation assays. **I**. Mammosphere formation assays of MB231 cells treated with ClpP agonists. Data shown as ave +/-SEM, summary of 3 independent experiments. *p*=0.017, One-way ANOVA. (p<0.05) Dunnett’s multiple comparisons test (compared with Ctl.). **J**. Representative images of mammosphere assays in culture for 14 days. **K**. Mammosphere formation assays with MB231 CLPP WT vs KO cell lines. Data shown as ave+/-SEM, summary of 3 independent experiments. \*\**p*<0.01, N.S.; not significant, Student’s *t*-test.

### ClpP agonists inhibit CSC function *in vitro*

To examine if ClpP agonists inhibit the fraction of CSCs, ALDEFLUOR assays were performed. The fraction of ALDEFLUOR bright (BR) cells was decreased from 0.34 % to 0.042 % (87.6 % reduction) in MB231 cells (Fig.2F) and from 7.64 % to 1.74 % (77.2 % reduction) in SKBR3 cells (Fig.S2G) after treatment with TR-57 for 48 h. The SORE6+ reporter assay was also used to examine the effect of ClpP agonist on CSCs. MB231 stably transfected with the lentiviral SORE6+ reporter gene were treated for different durations (24-72 h). We observed that ONC201 significantly reduced the SORE6+ fraction compared with DMSO control (Fig.2G).

Next, we determined the effects of ClpP agonists on mammosphere formation. Adherent cells were pretreated with drugs for 48 h, then equal number of viable cells were re-plated to low attachment chambers and incubated 10-14 days in the absence of drug (Fig.2H, I&J). ClpP agonists inhibited mammosphere formation in CLPP WT cells but not in CLPP KO cells (MB231, Fig.2I, J, K; MCF7 Fig.S2H; SUM159 Fig.S2I).

### ClpP agonists inhibit tumor initiation *in vivo*

Next, we assessed the impact of ClpP agonists on CSC *in vivo*. Similar to the mammosphere assays described above (Fig.2H), MB231 cells were pretreated with ONC201(5 μM) or DMSO. After 48 h, cells were trypsinized and collected, 5×10^5^ or 5×10^6^ live cells were injected into mouse MFP, and tumor formation was monitored (Fig.S3A). Tumor formation was detected 10 days after injection. DMSO-treated cells showed higher rate of tumor formation compared with ONC201-treated cells at both cell density groups (Fig.S3B). At Day 20, 90 % of mice injected with 5×10^6^ cells formed tumors in DMSO group, whereas only 40 % mice formed tumors in ONC201-group. One mouse injected with 5×10^5^ cells developed tumor in DMSO group, while no tumor formation was detected with ONC201-group. ELDA(19) indicated that CSC frequency was significantly (*p*<0.05) decreased by ONC201 at multiple time points (Fig.S3C). The average tumor volume was smaller in ONC201-treated cells compared with vehicle control group (Fig.S3D). To examine if tumors grown in mice injected with ONC201-treated cells were resistant to ONC201, 5 tumors from control group (5×10^6^ cells/mouse) and 5 tumors ONC201 group (5×10^6^ cells/mouse) were collected, and human cells were isolated from tumor tissue, and treated with ONC201. Cells harvested from the mouse tumors pretreated with ONC201 were still equally sensitive to ONC201 compared to the DMSO control (i.e., the 48 h treatment with ONC201 did not induce or select for resistance; Fig.S3E).

*In vivo* experiments were repeated comparing tumor formation by TR-57-treated or control MB231 CLPP WT and KO cells (Fig.3A). MB231 CLPP WT or CLPP KO cells were pretreated with TR-57 or DMSO 48 h, 5×10^5^ or 5×10^6^ cells live cells were injected into mouse MFP, and tumor formation was monitored. DMSO-treated CLPP WT cells developed tumors (30 % in 5×10^5^ group, 40 % in 5×10^6^ cells group) by Day 45, whereas TR-57-treated CLPP WT cells did not form tumors at either cell concentration (Fig.3B). By ELDA, CSC frequency was significantly inhibited (*p*=0.0007) in TR-57-treated CLPP WT cells compared with the DMSO - treated CLPP WT cells (Fig.3C). Importantly, no difference between control and TR-57 treated cells was seen in CLPP KO cells (Fig.3B&C). These findings further support that ClpP agonists impair breast CSC function in a CLPP-dependent fashion.

**Fig.3.**
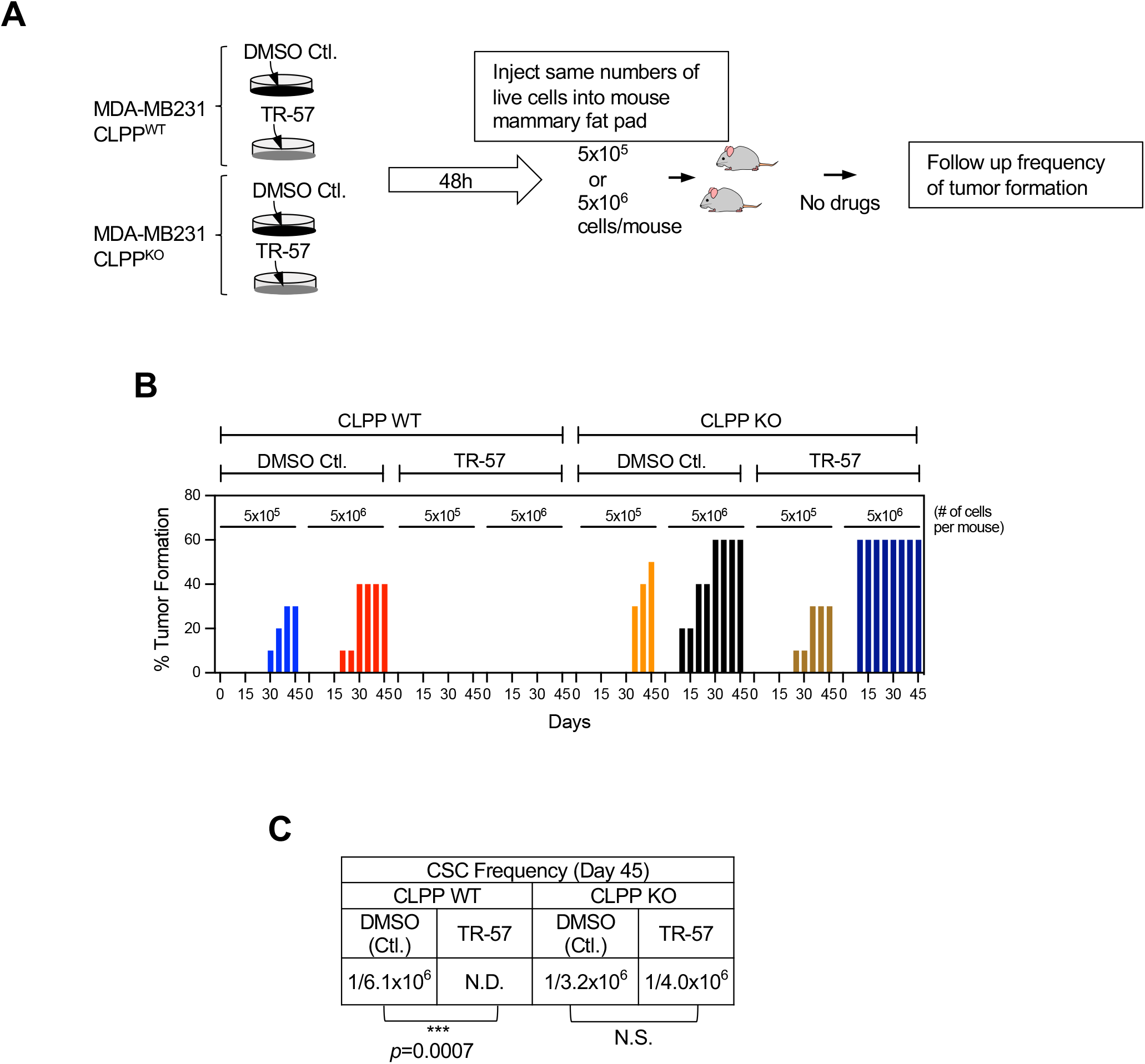
ClpP agonists inhibit tumor initiation *in vivo*. **A**. Experimental procedure illustrating 2^nd^ *in vivo*. **B**. Tumor formation (%) in each group at different time points up to Day 45. **C**. CSC Frequency between Ctl. and TR-57-treated groups, as well as CLPP WT and KO cells was determined using ELDA software.

### TR-57 is more effective at inhibiting mammosphere formation compared with other mitochondria targeting drugs

Previous studies have shown that other mitochondria-targeting drugs, such as oligomycin, metformin, and CPI-613, a pyruvate dehydrogenase/alpha-ketoglutarate dehydrogenase inhibitor decrease CSC function(20,21). We previously reported that oligomycin (IC_50_ ∼1-2 μM) and metformin (IC_50_ ∼10 mM) inhibit cellular ATP levels in MB231 cells(3). CPI-613 showed cytotoxicity in MB231 (IC_50_ ∼400 μM) and SUM159 (IC_50_ =126 μM, Fig.S4A).

We compared ClpP agonists and these drugs in mammosphere formation assays. Cells were pretreated with drugs at approximately their IC_50_ for 48 h, and then equal number of live cells were re-plated onto low attachment chambers without drugs (Fig.4A). TR-57 significantly inhibited mammosphere formation, oligomycin had a modest effect, while metformin and CPI-613 did not show an inhibitory effect. To test if the wash-out of drug at 48 h accounted for the differential sensitivities, we directly seeded cells into low attachment chambers, the drugs were added the next day and were left in the culture throughout the experiment (Fig.4B). In this setting, only TR-57 inhibited mammosphere formation. We also tested mammosphere assay formation with repeated drug administration every 2-3 days (Fig.4C). In this setting, all the drugs inhibited mammosphere formation but again TR-57 was most effective. These observations indicated that mitochondria-targeting drugs have a capacity to inhibit CSC functions in general. However, while the effects of the other drugs tested require repeated exposure to exert their inhibitory effect, ClpP agonists appear to require only a single short treatment (e.g, 48 h).

**Fig.4.**
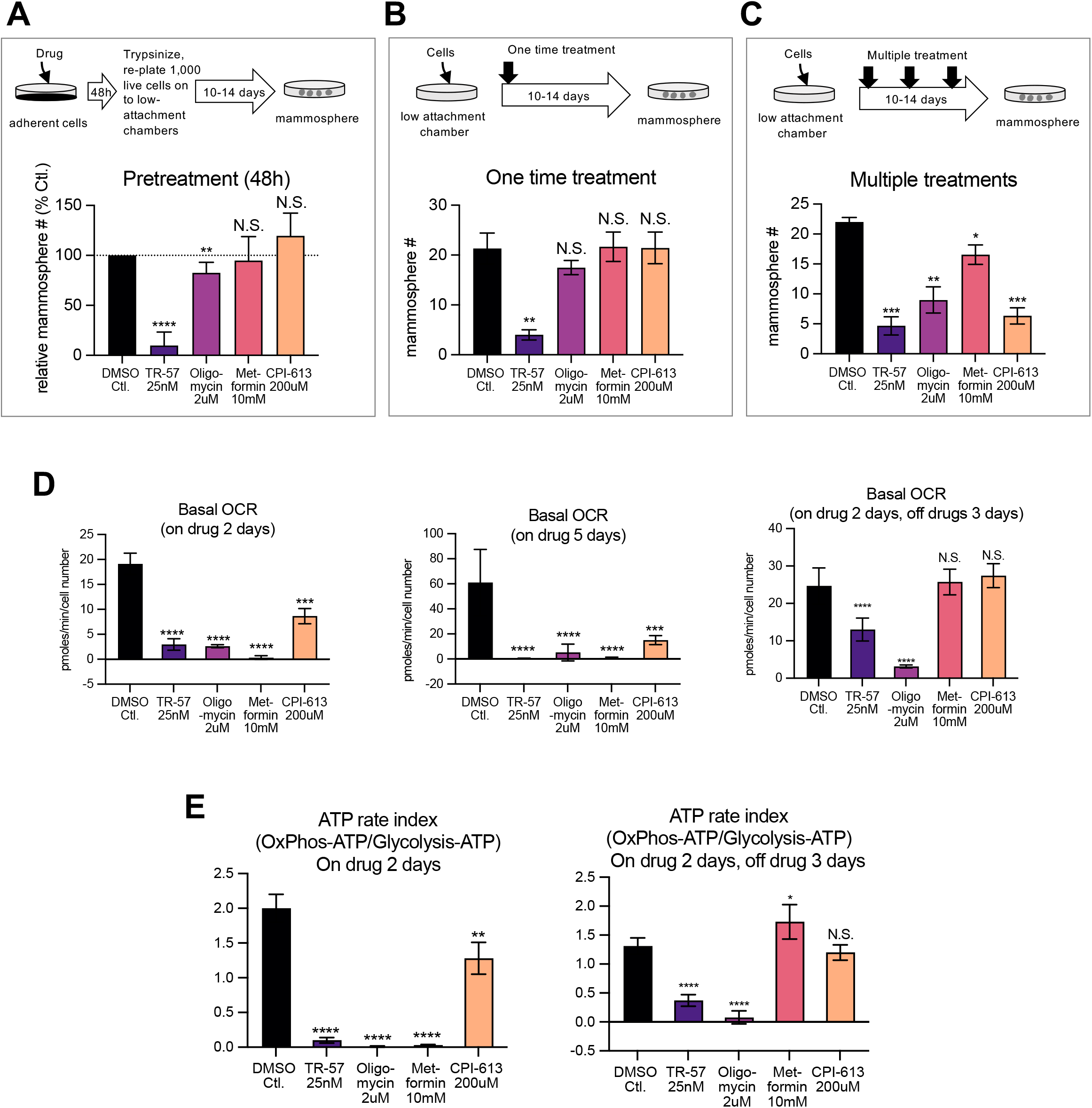
TR-57 exhibit higher potency to inhibit mammosphere formation compared with other mitochondria targeting drugs. **A**. Mammosphere formation assays with 2 days drug pretreatment. Experimental procedure (top) and results shown as ave+/-SD (bottom), summary of multiple (DMSO n=5, others n=3) experiments. **B**. Mammosphere formation assays with one time drug treatment. Experimental procedure (top) and results shown as ave+/-SD (bottom), summary of 3 independent experiments. **C**. Mammosphere formation assays with multiple dosing. Experimental procedure (top) and results shown as ave+/-SD (bottom), summary of 3 independent experiments. **D**. OCR measured with XF analyzer with multiple mitochondria-targeting drugs, different treatment durations. **E**. ATP rate index of cells treated with multiple mitochondria-targeting drugs, with or without drug washout. MB231 cells were used in all the experiments shown in Fig.4. \**p* <0.05, \*\**p*<0.01, \*\*\**p*<0.001, \*\*\*\**p*<0.0001, Student’s *t*-test.

To confirm the effects of the mitochondria-targeting drugs, basal OCR was measured at the IC_50_ for each. We observed that all the drugs significantly inhibited OCR, and the inhibitory effects remained even after 5 days of treatment without additional dosing (Fig.4D). When drugs were washed out after 48 h treatment, a significant inhibitory effect was still detected with TR-57 and oligomycin, while basal OCR of cells treated with metformin and CPI-613 were completely reversed upon drug washout (Fig.4D). This suggested that inhibitory effects of TR-57 and oligomycin on mitochondrial respiration are irreversible over the time frame examined, whereas that of metformin and CPI-613 are reversible after drug washout. Similarly, after 48 h of drug treatment, all the drugs significantly decreased OxPhos-ATP based on the ATP rate assay (Fig.4E, left). When drugs were removed at 48 h and ATP assay was examined 3 days post washout, the inhibitory effects of TR-57 and oligomycin remained, while that of metformin and CPI-613 was no longer detected (Fig.4E, right). Altogether, TR-57 most effectively inhibited mammosphere formation compared with other mitochondria-targeting drugs. In addition, while both TR-57 and oligomycin comparably inhibited mitochondrial respiration, TR-57 inhibited CSC function more efficiently, implying that the inhibitory effect of TR-57 on CSC is not solely dependent on OxPhos inhibition.

### ClpP agonists and other mitochondria-targeting drugs downregulate multiple pathways involved with maintenance of CSC

To investigate the mechanisms by which ClpP agonists inhibit CSC functions, we first interrogated RNAseq analysis of MB231 treated with ONC201 for 0, 3, 6, 12, and 24 h(3). GSEA indicated that transcripts involved with Myc target genes, cholesterol homeostasis were significantly downregulated by ONC201 at 24 h (Fig.S5A). IPA also suggested that the mevalonate pathway is significantly dysregulated by ONC201(Fig.S5B). Metacore enrichment analysis indicated that in addition to mevalonate pathway, HIF and YAP/TAZ pathways are dysregulated (Fig.S5C). Together, RNAseq analysis suggested that ONC201 dysregulates multiple signaling pathways and proteins critical for CSC maintenance, including the mevalonate pathway(22), HIF1α/HIF2α(23), YAP(24), and Myc(20,23,25).

We previously showed that ONC201 induced AMPK activation(3). Prior work has shown that AMPK phosphorylates YAP at Ser94 inhibiting YAP activity(26). We observed that TR-57 induces phosphorylation of YAP at Ser94 in a ClpP-dependent manner (Fig.S6A&B). Both ONC201 and TR-57 inhibited HOPflash reporter activity, an indicator of YAP/TAZ transcriptional activity (Fig.S6C). SiRNA mediated knockdown of YAP/TAZ impaired mammosphere formation (Fig.S6D&E), consistent with previous studies(27,28).

Cholesterol biosynthesis is also a key characteristic in breast CSCs(22) and is a positive regulator of YAP/TAZ activity(29). The mevalonate-YAP/TAZ axis is required for breast CSCs function(30). TR-57 downregulated 3-Hydroxy-3-Methylglutaryl-CoA Synthase 1 (HMGCS1), one of the critical enzymes in mevalonate pathway in a CLPP-dependent manner (Fig.S6F&G). In addition, simvastatin inhibited HOPflash reporter activity (Fig.S6H) and mammosphere formation (Fig.S6I), consistent with the reported link between the mevalonate pathway and YAP pathway, and the critical role of the mevalonate-YAP/TAZ axis in CSC functions. Similar to TR-57, simvastatin also induced AMPK activation and YAP phosphorylation at Ser94, and it was reversed by mevalonolactone (Fig.S6J), suggesting that inhibition of mevalonate pathway is correlated with impairment of YAP activity via AMPK activation. Together, we observed that a ClpP agonist downregulates mevalonate-YAP/TAZ axis, resulting in inhibition of CSC function. However, further analysis revealed that these findings are not specific to TR-57. Other mitochondria-target drugs also downregulated HMGCS1, HOPflash activity (Fig.S6L), oligomycin activated AMPK, and all increased phosphorylated YAP at Ser94 (Fig.S6M&N). These findings implied that mitochondria-targeting drugs downregulate YAP/TAZ and the mevalonate pathway as common targets, thereby leading to inhibition of breast CSC function.

We observed that TR-57 downregulated Myc at the protein level in a CLPP-dependent manner (Fig.S7A) but did not inhibit the transcript level (Fig.S7B). While TR-57 decreased Myc phosphorylation at Ser62 which stabilizes Myc, TR-57 transiently induced phosphorylation of Myc at Thr58 (Fig.S7C), which leads to proteasomal degradation of Myc(31). Consistent with previous studies(20,25), siRNA-mediated suppression of Myc inhibited mammosphere formation (Fig.S7D&E). The other mitochondria-targeting drugs also downregulate Myc (Fig.S7F). Myc is one of the target genes of YAP/TAZ pathway(32), and indeed suppression of YAP/TAZ decreased Myc (Fig.S7G&H).

Western blot revealed that HIF1α was mostly detected in the nuclear fraction and total and hydroxylated HIF1*α* were transiently increased upon TR-57 treatment follow by a marked decrease (Fig.S7I). Similarly, HIF2α appeared to be downregulated (Fig.S7I), while transcripts of HIF1α and HIF2α (gene name: EPAS1) were not decreased by TR-57 (Fig.S7J). SiRNA-mediated knockdown of EPAS1 and dual knockdown of HIF1α and EPAS1 impaired mammosphere formation (Fig.S7K&L). HIF transcriptional activity measured with HRE-Luc reporter gene was downregulated by ClpP agonists, oligomycin, and metformin in both MB231 and SUM159 cells (Fig.S7M).

In summary, we observed that multiple pathways critical for CSC function, such as the YAP pathway, mevalonate pathway, Myc, HIF1α/HIF2α pathways, are dysregulated by mitochondria-targeting drugs, but these effects are not specific to ClpP agonist.

### ClpP agonists dysregulate pathways not affected by other mitochondrial drugs

#### ClpP agonist downregulates NAD(P)+ and NAD(P)H and induces oxidative stress

We investigated other mitochondrial pathways that might be inhibited uniquely or more effectively by ClpP agonists. Recent studies have demonstrated essential roles of NAD(P)+ on stem cell pluripotency(33) and NAD(P)H is considered a CSC marker(34). Both NAD(P)+ and NAD(P)H are directly involved in redox homeostasis(35,36), which is pivotal to maintain self-renewal capacity of stem cells(37). Therefore, we examined the impact of TR-57 on these co-factors and redox homeostasis. TR-57 decreased total levels of NAD+ and NADH and NADP+ and NADPH in a CLPP-dependent manner (Fig.5A&B, MB231 top, SUM159 bottom).

**Fig.5.**
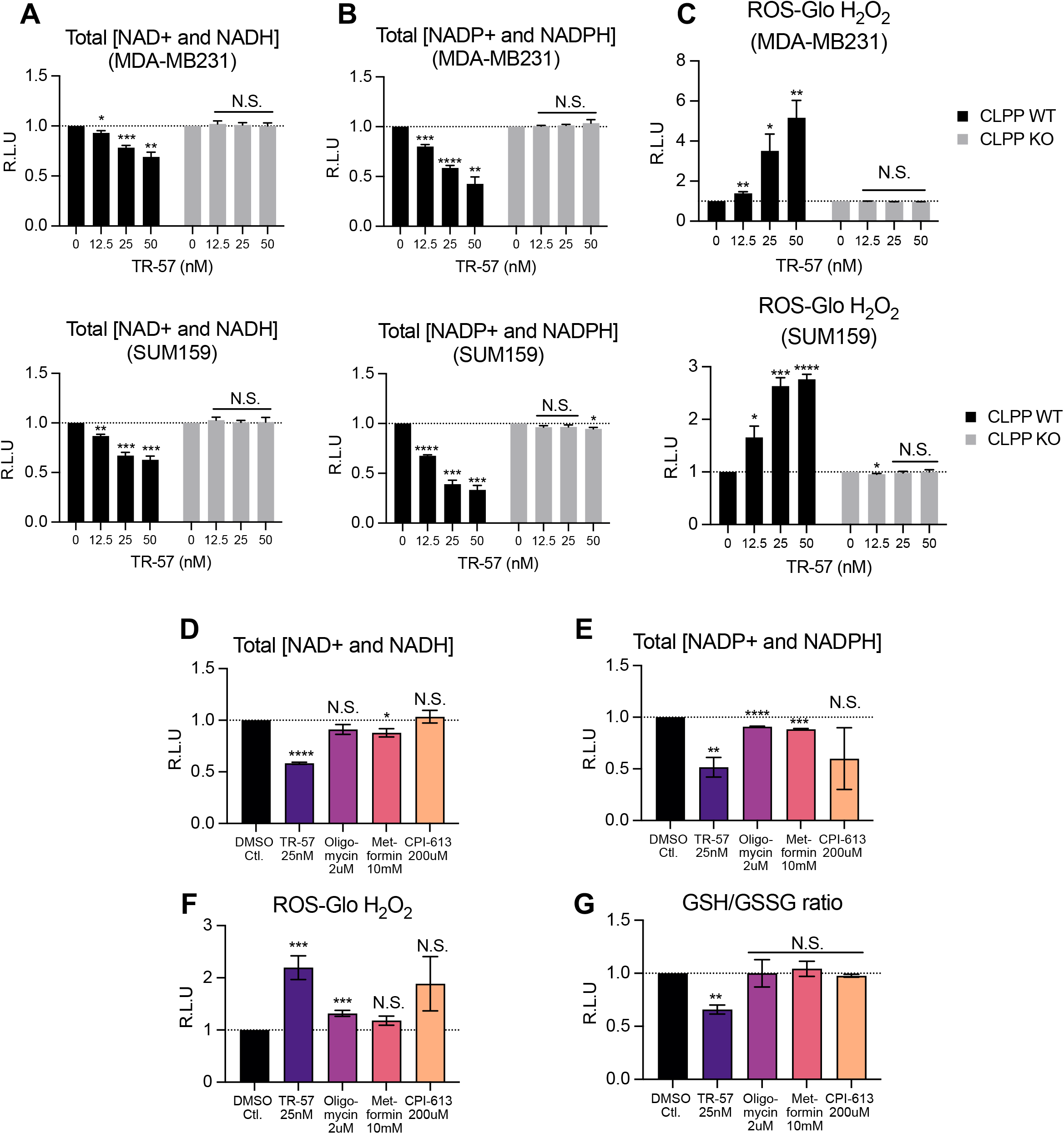
ClpP agonist downregulates NAD(P)/NAD(P)H and induces oxidative stress. **A**. total level of NAD+/NADH, **B**. total level of NADP+/NADPH, and **C**. ROS-Glo H_2_O_2_ assays with MB231 (top) and SUM159 (bottom) cells treated with TR-57 for 5 days. Data shown as ave+/SEM, summary of 3 independent experiments. **D**. Total level of NAD+/NADH assays (n=3) **E**. Total NADP+/NADPH assays (n=3), **F**. ROS-Glo H_2_O_2_ assays (n=5), **G**. GSH/GSSG ratio (n=3), MB231 cells treated with multiple mitochondria-targeting drugs for 5 days. Data shown as ave+/- SEM, summary of multiple experiments as indicated.

Additionally, TR-57 significantly elevated ROS in a CLPP-dependent manner (Fig.5C, MB231 top, SUM159 bottom). Compared with TR-57, other mitochondria-targeting drugs showed less or no inhibitory effects on total level of NAD+ and NADH, or NADP+ and NADPH (Fig.5D&E). TR-57 induced the largest increase in ROS and decreased the GSH/GSSG ratio compared to the other mitochondria targeting drugs (Fig.5F&G). Together, these observations indicated that ClpP agonist more profoundly dysregulates NAD+ and NADH, NADP+ and NADPH, and impairs redox homeostasis compared to the other drugs.

Next, we tested if the decreased levels of NAD+ and NADH account for some of TR-57 effect on cell viability and CSC function. FK866, a nicotinamide phosphoribosyltransferase (NAMPT) inhibitor, was used as a positive control of NAD+ and NADH depletion. As expected, FK866 significantly impaired cell viability in MB231 cells (IC_50_=1.93 nM), depleted total NAD+ and NADH, increased ROS (Fig.S8A), and inhibited mammosphere formation (Fig.S8B), as previously reported (38). The inhibitory effect of FK866 on cell viability and CSC function was specifically due to lack of NAD+, as it was completely reversed by two supplemental nicotinamides (nicotinamide riboside [NR] and nicotinamide mononucleotide [NMN]) (Fig.S8C&E). In contrast, the effect of TR-57 was not rescued by these nicotinamides (Fig.S8D&E). This confirmed that NAD+ is required for CSC function in accordance with other reports(33,38), however, also indicated that NAD depletion is not the sole mechanism by which ClpP agonists impair CSC. In contrast to nicotinamides, N-acetyl-cysteine (NAC), a ROS scavenger, did not reverse cell viability and mammosphere formation inhibited by FK866 and TR-57.

#### ClpP agonist downregulates multiple mitochondrial enzymes involved with folate-mediated one carbon metabolism (FOCM)

Cellular NADPH is largely generated by the pentose phosphate pathway (PPP), FOCM, and malic enzymes (ME) in cancer and proliferating cells(35) (Fig.S9). Our finding that TR-57 depletes NADP+ and NADPH (Fig.5B) suggested that TR-57 downregulates those pathways/enzymes.

We observed that TR-57 downregulates multiple enzymes involved with FOCM, including mitochondrial one-carbon metabolism enzyme methylene tetrahydrofolate dehydrogenases 2 (MTHFD2), serine hydroxymethyltransferase 2 (SHMT2), thymidine synthase (TYMS), and malic enzyme 2 (ME2), in both dose, time, and CLPP-dependent manner (MB231 Fig.6A&B, SUM159 Fig.S10A&B). Importantly, downregulation of these enzymes was a ClpP-agonist specific effect, and it was not observed by other mitochondria-targeting drugs (Fig.S10C). Conversely, D-3-phosphoglycerate dehydrogenase (PHGDH) which mediates serine biosynthesis, and cystathione gamma-lyase (CGL/CTH), an enzyme involved with one-carbon metabolism outside of mitochondria (Fig.S9), were increased by TR-57 (MB231, Fig.6A&B, Fig.S10C, SUM159 Fig.S10A&B). Similarly, glucose-6-phosphate-dehydrogenase (G6PD), a rate-limiting enzyme of the PPP (Fig.S9), was also slightly increased by TR-57 (MB231, Fig.6A&B). Moreover, enzymatic activity of G6PD was increased by TR-57 (Fig.S10D). These increases are likely compensatory changes.

**Fig.6.**
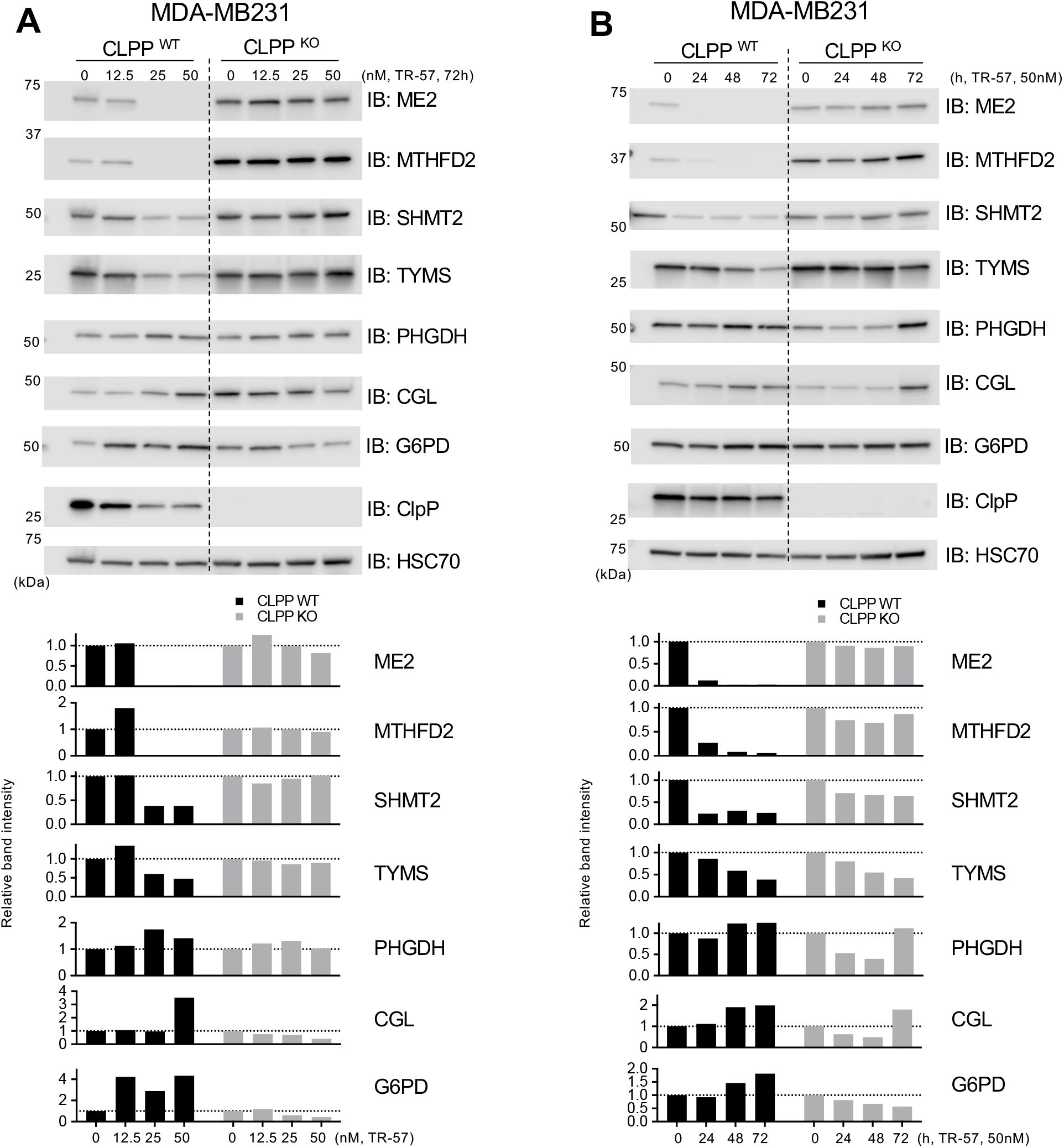
ClpP agonist inhibits folate mediated one carbon metabolism. **A&B**. Immunoblots showing dose (**A**) and time (**B**) dependent effects of TR-57 on enzymes involved with one-carbon metabolism, serine synthesis pathway, and PPP in MB231 CLPP WT and KO cell lines. Representative data from multiple experiments are shown. Relative band intensities of each protein are shown in lower panels.

#### ClpP agonist downregulates multiple mitochondrial enzymes involved with glutamine-proline axis and impairs proline biosynthesis

Proteomics analysis (Emily M.J. Fennell, unpublished data) indicated that ClpP agonists significantly decrease levels of glutaminase/GLS, ALDH18A1, pyrroline-5-carboxylate reductase 1 and 2 (PYCR1/2), enzymes involved with the glutamine-proline axis (Fig.7A). GLS converts glutamine to glutamate in mitochondria, glutamate is converted to pyrroline-5-carboxylate (P5C) via ALDH18A1. P5C is further reduced to proline by PYCR1/2(39). PYCR1/2 are essential mitochondrial enzymes for proline biosynthesis(40).

**Fig.7.**
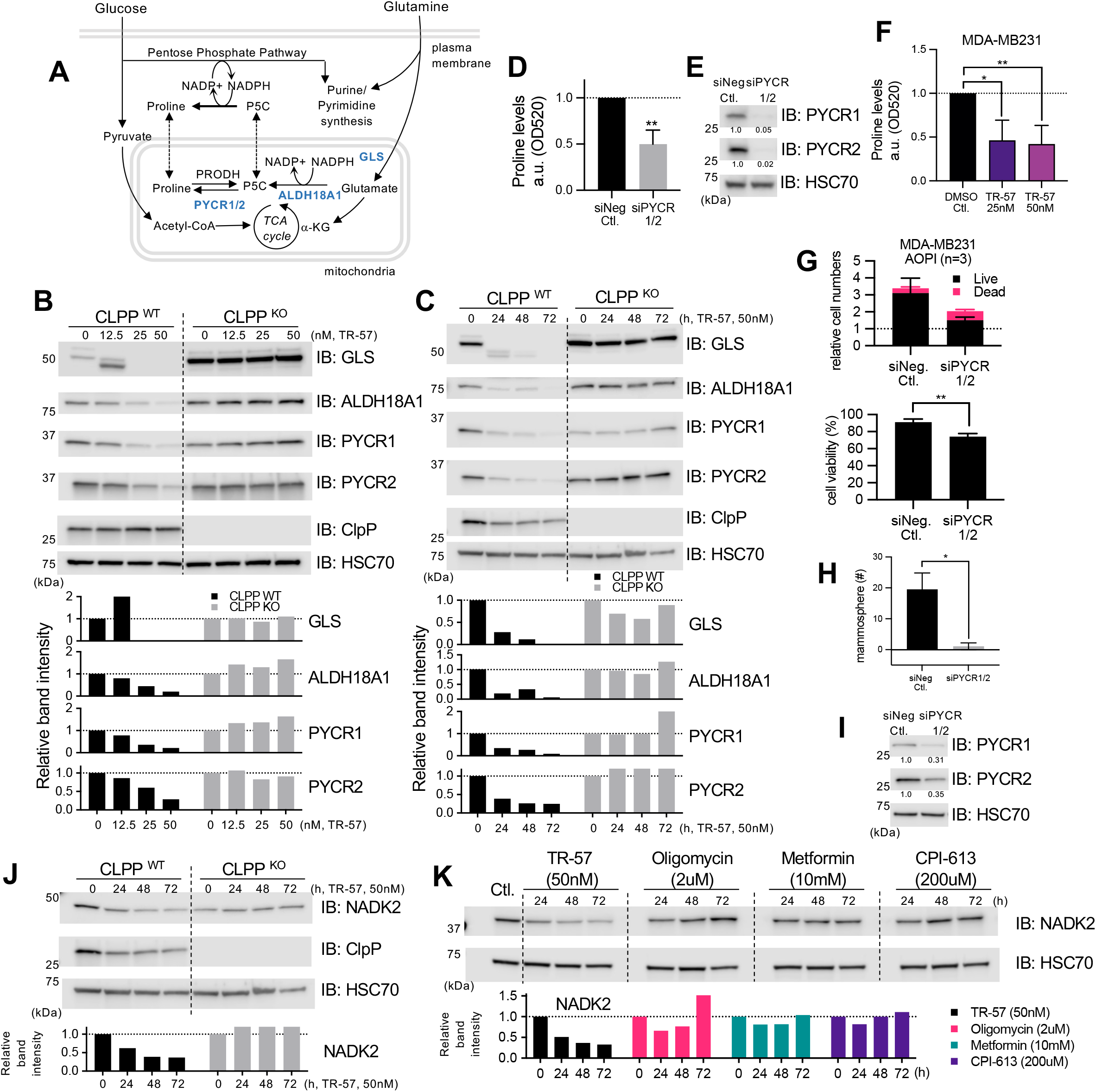
ClpP agonist downregulates proline biosynthesis. **A** Diagram illustrating glutamine-proline axis and mitochondrial enzymes involved. **B&C**. The dose(**B**) and time(**C**)-dependent effect of TR-57 on enzymes involved with glutamine-proline axis in MB231 cells. Relative band intensities of each protein are shown in lower panels. **D**. Proline assays. Intracellular proline levels were measured with SUM159 cells transfected with control siRNA or PYCR1/2 siRNA, 3 days after siRNA transfection. Data shown as ave+/-SD, summary of 3 independent experiments. **E**. Representative immunoblots showing knockdown of PYCR1/2 in SUM159. **F**. Proline assays in MB231 cells treated with DMSO or TR-57 for 72h. Data shown as ave+/-SD, summary of 3 independent experiments. \**p*<0.05, \*\**p*<0.01, Student’s *t*-test. **G**. Relative cell viabilities analyzed with AOPI assays. Cell viability of MB231 was examined after 3 days of transfection with control siRNA or PYCR1/2 siRNA. Data shown as ave+/-SD, 3 independent experiments. Top: cell numbers of live and dead cells relative to initial cell numbers transfected. Bottom: % of cell viability (live/total cell numbers). \*\**p*<0.01, Student’s *t*-test. **H**. Mammosphere formation assays with MB231 cells transfected with control siRNA or siPYCR1/2. Data shown as ave+/-SEM, summary of 3 independent experiments. \**p*<0.05, Student’s *t*-test. **I**. Western blot showing knockdown of PYCR1/2 by siRNA, 48h after transfection. **J**. Time-dependent effect of TR-57 on NADK2 in MB231 CLPP WT and KO cell lines. Relative band intensity of NADK2 is shown in lower panel. **K**. Time-dependent effect of multiple mitochondria-targeting drugs on NADK2 in MB231. Relative band intensity of NADK2 is shown in lower panel.

Western blotting confirmed that GLS, ALDH18A1, PYCR1/2 were all downregulated by TR-57 in MB231 (Fig.7B&C) and SUM159 (Fig.S11A&B) in a time, dose, and CLPP-dependent manner. Other mitochondria-targeting drugs caused some downregulation of GLS and ALDH18A1 but did not downregulate PYCR1/2 (Fig.S11C).

Next, we investigated the role of PYCR1/2 in proline biosynthesis in breast cancer cells. Knockdown of PYCR1/2 impaired proline biosynthesis (Fig.7D&E). Similarly, TR-57 inhibited proline biosynthesis (MB231: Fig.7F, SUM159: Fig.S11D), presumably reflecting the downregulation of PYCR1/2 protein (Fig.7B&C, S11A&B). PYCR1/2 knockdown significantly inhibited cell growth (Fig.7G in MB231, Fig.S11E&F in SUM159), consistent with a previous report(40). Furthermore, we found that PYCR1/2 knockdown significantly inhibited mammosphere formation (Fig.7H&I). This suggested that proline biosynthesis is required for cell growth and CSC function in breast cancer cells. In contrast to the effects of PYCR1/2 loss, inhibition of GLS with CB-839 only modestly inhibited cell viability but did not impair mammosphere formation (Fig.S11G&H). The ROS level induced by CB-839 was not as high as that of TR-57, and GSH/GSSH ratio was not changed by CB-839 (Fig.S11I).

Two recent studies reported that mitochondrial NADP(H) is essential for proline biosynthesis during cell growth, and that mitochondrial NAD kinase 2 (NADK2), an enzyme responsible for production of mitochondrial NADP^+^, is vital for proline biosynthesis(41,42)(Fig.S9). We observed that NADK2 is also reduced by TR-57 in a CLPP-dependent manner (Fig.7J for MB231, Fig.S11J&K for SUM159), and this was not observed by other mitochondria-targeting drugs (Fig.7K). In summary, we show that loss of PYCR1/2 impairs proline biosynthesis and inhibits mammosphere formation. TR-57 but not the other mitochondrial targeting drugs, impairs proline biosynthesis by targeting multiple enzymes involved with glutamine-proline axis, contributing to growth inhibition and CSC inhibition.

### Expression of mitochondrial enzymes involved with glutamine-proline axis and FOCM in breast cancer cell lines and patients samples with different molecular subtypes

RNAseq analysis of 15 breast cancer cell lines indicated that there was a trend to higher expression of GLS in TNBC cell lines compared with ER+ and HER2 amplified cell lines but not of other genes in these metabolic pathways (Fig.S12A). Western blot of 17 breast cancer cell lines showed higher protein expression of GLS and TYMS in TNBC cell lines compared to ER+ and HER2 amplified cell lines, and higher expression ALDH18A and PYCR2 in TNBC compared to ER+ cell lines. HER2 amplified cell lines showed higher TYMS and ALDH18A1 compared with ER+ cell lines (Fig.S12B&C). Proteomic profiling of small number of breast cancer patients with different subtypes (43) indicated that GLS and TYMS are highest in TNBC (Fig.S12D), consistent with the immunoblot data from cell lines (Fig.S12B&C). ALDH18A1 was increased in HER2 amplified and TNBC, compared to luminal breast cancers, again consistent with the cell line data but was highest in the HER2 amplified tumors. The data for PYCR2 did not show an increase in TNBC (Fig.S12D), which differed from the cell line data.

## Discussion

We previously reported that ONC201 disrupts mitochondria structure and function in breast cancer cells(3). In the present study, we further investigated the impact of ONC201 and the more potent TR ClpP agonists in breast CSC function. We confirmed that the mechanism of action of ONC201 and TR-compounds is dependent on CLPP using CLPP KO cells. Further, we show that ClpP agonists inhibited CSC function *in vitro* and *in vivo* in a CLPP-dependent fashion. ClpP agonist showed greater inhibitory effect on mammosphere formation compared with other mitochondria-targeting drugs, despite the comparable inhibition of OxPhos. We found that ClpP agonists and other mitochondria-targeting drugs tested downregulate multiple CSC signaling pathways such as YAP, the mevalonate pathway, Myc, and HIF. We also found that ClpP agonists uniquely dysregulate additional mitochondrial metabolic pathways critical to CSC function. ClpP agonists significantly deplete NAD(P)+ and NAD(P)H and dysregulate redox homeostasis. ClpP agonists downregulate multiple NADPH generating enzymes involved with FOCM. Moreover, we observed that ClpP agonists inhibit glutamine-proline axis and found that proline biosynthesis is critical for breast CSC function. Thus, we report that ClpP agonists widely dysregulate mitochondrial functions, including bioenergetic, biosynthetic, and signaling pathways, leading to inhibition of CSCs function (Fig.8).

**Fig.8.**
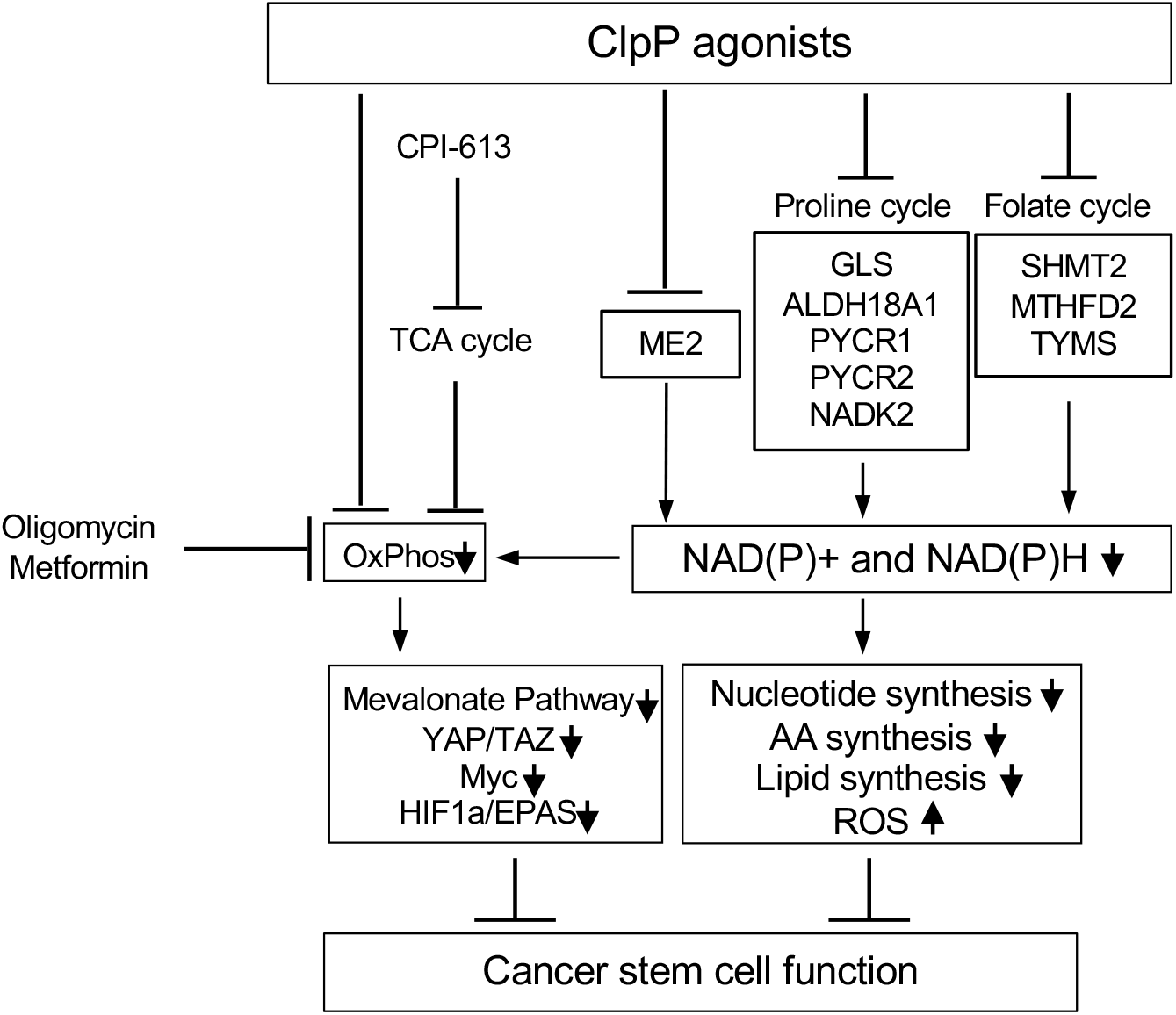
Graphical summary of mechanism of action of ClpP agonist in cancer cells. ClpP agonists inhibit breast cancer cell growth and CSC functions by disrupting mitochondrial metabolic homeostasis, in addition to OxPhos inhibition. Other mitochondrial targeted drugs have more restricted effects on mitochondria as indicated.

Prior work has shown that drugs that inhibit OxPhos impair CSC function but the mechanisms responsible for this have not be clearly identified(44). Besides the metabolic dysregulation, we observed that multiple CSC signaling pathways were downregulated by all of the mitochondria-targeting agents tested (Fig.S6, S7). We found a complex network of inter-connected CSC pathways that are downregulated by ClpP agonists as well as other mitochondrial inhibitors (Fig.S13). While ATP depletion may directly downregulate mevalonate pathway(45), our findings suggest that activated AMPK plays an important role mediating ATP depletion and downregulation of mevalonate pathway and YAP pathway. Prior studies have shown that activated AMPK inhibits the mevalonate pathway(46), and conversely, inhibition of mevalonate pathway activates AMPK(47). A previous study showed that cellular energy stress induces AMPK-mediated inhibition of YAP by phosphorylation at Ser94(26). We found that ClpP agonists and other mitochondria-targeting drugs activated AMPK, leading to inhibition of YAP via phosphorylation at Ser94. Our finding is the first to show the link between OxPhos inhibition and YAP Ser94 phosphorylation. ClpP agonists and other mitochondria-targeting drugs downregulated Myc, and this is also shown in recent reports(48-50). Myc stimulates mitochondrial biosynthesis, including glutamine-proline axis(51-53), therefore downregulation of Myc by ClpP agonists may lead to further mitochondrial dysfunction(39,52). We observed that the HIF pathway was also downregulated by OxPhos inhibitions, although the mechanism remains unclear. Possibly, OxPhos inhibitors induce accumulation of intracellular O_2_, leading to HIF1α/HIF2α destabilization(54). Additionally, HIF pathway is regulated by AMPK(55) and Myc(56) (Fig.S13).

The second novel finding from our work is that ClpP agonists, but not the other mitochondria-targeting agents, downregulated multiple enzymes involved with glutamine-proline axis. Glutamine catabolism has been shown to be critical for inhibiting breast CSC function(57). We demonstrated for the first time that PYCR1/2, but not GLS, is critical for breast CSC function. Indeed, higher PYCR1 mRNA levels were significantly associated with poor survival in breast cancers, regardless of ER status(58). Recent studies have shown that proline metabolism plays multiple roles in cancer, such as nucleotide/protein synthesis, generation of NAD+, redox homeostasis(40,51,59). Thus, proline metabolism is proposed as a promising therapeutic target in breast cancer(39,59,60) and our findings extend the importance of proline by demonstrating a role in CSC function.

We also found that ClpP agonists downregulate multiple NADPH generating enzymes such as ME2, MTHFD2, SHMT2, TYMS, all of which have been linked to CSC function(61-64)(Fig.6, Fig.S10). NADPH provides strong reducing power and maintains redox homeostasis in a cell, and this is particularly crucial for CSCs to maintain the low ROS levels(65). Targeting NADPH has been considered as a rational strategy for cancer treatment(66), however, it is challenging because NADPH metabolism is shared in normal and cancer cells(35). With respect to this selectivity, as we previously reported, ClpP agonists do not impair cell viability in non-transformed cells(3). Why non-transformed cells are resistant to ClpP agonists remains to be addressed. Nevertheless, our results suggest that modulating NADPH by ClpP agonist contributes to inhibiting breast CSC function.

The ClpP agonists inhibited CSC function *in vitro* and *in vivo* (Fig.2&3, Fig.S2&S3), suggesting that ClpP agonists may be effective in preventing metastasis and further studies are required to examine the efficacy of ClpP agonists on CSCs using cancer metastasis or adjuvant models. Eradicating CSCs by mitochondria-targeting agents has been attempted in previous studies(13,67). Considering the intertumoral metabolic heterogeneity and plasticity, targeting CSCs with a single mitochondria-targeting agent may not be sufficient. To overcome this, targeting major metabolic pathways with combination therapies should be considered(44). In addition to development of combination therapies, stratification of patients will be necessary to achieve successful results in clinic. Glutamine-dependency substantially varies among breast cancer subtypes(68), and TNBC appear to be more dependent on FOCM and glutamine-proline axis (69,70)(Fig.S12). Additionally, identification of pharmacodynamic biomarkers of ClpP agonist activity is critical to monitor the drug on target activity. Profiling metabolic activity of metabolic changes induced by ClpP agonists may help identify patients who may benefit from ClpP agonists and identify pharmacodynamic biomarkers of activity(5).

ONC201 has been tested in multiple clinical trials and was well tolerated(71). In the last decade, multiple ClpP agonists have been developed(72). TR compounds have significantly higher potency compared with ONC201 and are similar to the IC50s of ONC206 and ONC212 as shown in the present study and previous publications(4,73). Further studies are necessary to identify the best-in-class ClpP agonist.

In conclusion, ClpP agonists are a novel category of mitochondria-targeting drugs which cause pleiotropic disruption of mitochondrial functions and CSC function. ClpP agonists are promising new anti-tumor drugs in breast cancers worth further preclinical and clinical evaluation.

## Acknowledgement

Authors are grateful to Chimerix Inc. (formerly Oncoceutics, Inc), for generously providing ONC201, Mr. Franklin Ning and Dr. Massimo Fantini (Women’s Malignancies Branch, CCR, NCI), Dr. William Telford (Experimental Transplantation and Immunotherapy Branch, CCR, NCI) for technical assistance of flow cytometry experiments, Dr. Ronald Gress (Experimental Transplantation and Immunotherapy Branch, CCR, NCI) for access to a Seahorse XFe24 analyzer. We also would like to thank members of the Lipkowitz lab for their discussion and support.

## References

1. Medini H, Cohen T, Mishmar D. Mitochondria Are Fundamental for the Emergence of Metazoans: On Metabolism, Genomic Regulation, and the Birth of Complex Organisms. Annu Rev Genet 2020;54:151–66

2. Li Y, Li Z. Potential Mechanism Underlying the Role of Mitochondria in Breast Cancer Drug Resistance and Its Related Treatment Prospects. Front Oncol 2021;11:629614

3. Greer YE, Porat-Shliom N, Nagashima K, Stuelten C, Crooks D, Koparde VN, et al. ONC201 kills breast cancer cells in vitro by targeting mitochondria. Oncotarget 2018;9:18454–79

4. Graves PR, Aponte-Collazo LJ, Fennell EMJ, Graves AC, Hale AE, Dicheva N, et al. Mitochondrial Protease ClpP is a Target for the Anticancer Compounds ONC201 and Related Analogues. ACS Chem Biol 2019;14:1020–9

5. Ishizawa J, Zarabi SF, Davis RE, Halgas O, Nii T, Jitkova Y, et al. Mitochondrial ClpP-Mediated Proteolysis Induces Selective Cancer Cell Lethality. Cancer Cell 2019;35:721–37 e9

6. Nouri K, Feng Y, Schimmer AD. Mitochondrial ClpP serine protease-biological function and emerging target for cancer therapy. Cell Death Dis 2020;11:841

7. Mabanglo MF, Bhandari V, Houry WA. Substrates and interactors of the ClpP protease in the mitochondria. Curr Opin Chem Biol 2021

8. Zhou HM, Zhang JG, Zhang X, Li Q. Targeting cancer stem cells for reversing therapy resistance: mechanism, signaling, and prospective agents. Signal Transduct Target Ther 2021;6:62

9. Loureiro R, Mesquita KA, Magalhaes-Novais S, Oliveira PJ, Vega-Naredo I. Mitochondrial biology in cancer stem cells. Semin Cancer Biol 2017;47:18–28

10. Vlashi E, Lagadec C, Vergnes L, Reue K, Frohnen P, Chan M, et al. Metabolic differences in breast cancer stem cells and differentiated progeny. Breast Cancer Res Treat 2014;146:525–34

11. Walsh HR, Cruickshank BM, Brown JM, Marcato P. The Flick of a Switch: Conferring Survival Advantage to Breast Cancer Stem Cells Through Metabolic Plasticity. Front Oncol 2019;9:753

12. Jones CL, Inguva A, Jordan CT. Targeting Energy Metabolism in Cancer Stem Cells: Progress and Challenges in Leukemia and Solid Tumors. Cell Stem Cell 2021;28:378–93

13. De Francesco EM, Sotgia F, Lisanti MP. Cancer stem cells (CSCs): metabolic strategies for their identification and eradication. Biochem J 2018;475:1611–34

14. Tang B, Raviv A, Esposito D, Flanders KC, Daniel C, Nghiem BT, et al. A flexible reporter system for direct observation and isolation of cancer stem cells. Stem Cell Reports 2015;4:155–69

15. Shaw FL, Harrison H, Spence K, Ablett MP, Simões BM, Farnie G, et al. A detailed mammosphere assay protocol for the quantification of breast stem cell activity. J Mammary Gland Biol Neoplasia 2012;17:111–7

16. Chari R, Yeo NC, Chavez A, Church GM. sgRNA Scorer 2.0: A Species-Independent Model To Predict CRISPR/Cas9 Activity. ACS Synth Biol 2017;6:902–4

17. Sanjana NE, Shalem O, Zhang F. Improved vectors and genome-wide libraries for CRISPR screening. Nat Methods 2014;11:783–4

18. Nabbi A, Riabowol K. Rapid Isolation of Nuclei from Cells In Vitro. Cold Spring Harb Protoc 2015;2015:769–72

19. Hu Y, Smyth GK. ELDA: extreme limiting dilution analysis for comparing depleted and enriched populations in stem cell and other assays. J Immunol Methods 2009;347:70–8

20. Lee KM, Giltnane JM, Balko JM, Schwarz LJ, Guerrero-Zotano AL, Hutchinson KE, et al. MYC and MCL1 Cooperatively Promote Chemotherapy-Resistant Breast Cancer Stem Cells via Regulation of Mitochondrial Oxidative Phosphorylation. Cell Metab 2017;26:633–47 e7

21. Bellio C, DiGloria C, Spriggs DR, Foster R, Growdon WB, Rueda BR. The Metabolic Inhibitor CPI-613 Negates Treatment Enrichment of Ovarian Cancer Stem Cells. Cancers (Basel) 2019;11

22. Ehmsen S, Pedersen MH, Wang G, Terp MG, Arslanagic A, Hood BL, et al. Increased Cholesterol Biosynthesis Is a Key Characteristic of Breast Cancer Stem Cells Influencing Patient Outcome. Cell Rep 2019;27:3927–38 e6

23. Yang L, Shi P, Zhao G, Xu J, Peng W, Zhang J, et al. Targeting cancer stem cell pathways for cancer therapy. Signal Transduct Target Ther 2020;5:8

24. Zanconato F, Cordenonsi M, Piccolo S. YAP/TAZ at the Roots of Cancer. Cancer Cell 2016;29:783–803

25. Yang A, Qin S, Schulte BA, Ethier SP, Tew KD, Wang GY. MYC Inhibition Depletes Cancer Stem-like Cells in Triple-Negative Breast Cancer. Cancer Res 2017;77:6641–50

26. Mo JS, Meng Z, Kim YC, Park HW, Hansen CG, Kim S, et al. Cellular energy stress induces AMPK-mediated regulation of YAP and the Hippo pathway. Nat Cell Biol 2015;17:500–10

27. Sorrentino G, Ruggeri N, Zannini A, Ingallina E, Bertolio R, Marotta C, et al. Glucocorticoid receptor signalling activates YAP in breast cancer. Nat Commun 2017;8:14073

28. Cordenonsi M, Zanconato F, Azzolin L, Forcato M, Rosato A, Frasson C, et al. The Hippo transducer TAZ confers cancer stem cell-related traits on breast cancer cells. Cell 2011;147:759–72

29. Santinon G, Pocaterra A, Dupont S. Control of YAP/TAZ Activity by Metabolic and Nutrient-Sensing Pathways. Trends Cell Biol 2016;26:289–99

30. Sorrentino G, Ruggeri N, Specchia V, Cordenonsi M, Mano M, Dupont S, et al. Metabolic control of YAP and TAZ by the mevalonate pathway. Nat Cell Biol 2014;16:357–66

31. Sears RC. The life cycle of C-myc: from synthesis to degradation. Cell Cycle 2004;3:1133–7

32. Yamaguchi H, Taouk GM. A Potential Role of YAP/TAZ in the Interplay Between Metastasis and Metabolic Alterations. Front Oncol 2020;10:928

33. Navas LE, Carnero A. NAD(+) metabolism, stemness, the immune response, and cancer. Signal Transduct Target Ther 2021;6:2

34. Yuan Y, Yan Z, Miao J, Cai R, Zhang M, Wang Y, et al. Autofluorescence of NADH is a new biomarker for sorting and characterizing cancer stem cells in human glioma. Stem Cell Res Ther 2019;10:330

35. Ju HQ, Lin JF, Tian T, Xie D, Xu RH. NADPH homeostasis in cancer: functions, mechanisms and therapeutic implications. Signal Transduct Target Ther 2020;5:231

36. Panieri E, Santoro MM. ROS homeostasis and metabolism: a dangerous liason in cancer cells. Cell Death Dis 2016;7:e2253

37. Chakrabarty RP, Chandel NS. Mitochondria as Signaling Organelles Control Mammalian Stem Cell Fate. Cell Stem Cell 2021;28:394–408

38. Bonuccelli G, De Francesco EM, de Boer R, Tanowitz HB, Lisanti MP. NADH autofluorescence, a new metabolic biomarker for cancer stem cells: Identification of Vitamin C and CAPE as natural products targeting “stemness”. Oncotarget 2017;8:20667–78

39. Tanner JJ, Fendt SM, Becker DF. The Proline Cycle As a Potential Cancer Therapy Target. Biochemistry 2018;57:3433–44

40. Liu W, Hancock CN, Fischer JW, Harman M, Phang JM. Proline biosynthesis augments tumor cell growth and aerobic glycolysis: involvement of pyridine nucleotides. Sci Rep 2015;5:17206

41. Zhu J, Schwörer S, Berisa M, Kyung YJ, Ryu KW, Yi J, et al. Mitochondrial NADP(H) generation is essential for proline biosynthesis. Science 2021;372:968–72

42. Tran DH, Kesavan R, Rion H, Soflaee MH, Solmonson A, Bezwada D, et al. Mitochondrial NADP. Nat Metab 2021;3:571–85

43. Tyanova S, Albrechtsen R, Kronqvist P, Cox J, Mann M, Geiger T. Proteomic maps of breast cancer subtypes. Nat Commun 2016;7:10259

44. Jagust P, de Luxán-Delgado B, Parejo-Alonso B, Sancho P. Metabolism-Based Therapeutic Strategies Targeting Cancer Stem Cells. Front Pharmacol 2019;10:203

45. Kumari A. Chapter 7 -Cholesterol Synthesis. xIn: Kumari A, editor. Sweet Biochemistry: Academic Press; 2018. p 27–31.

46. Motoshima H, Goldstein BJ, Igata M, Araki E. AMPK and cell proliferation--AMPK as a therapeutic target for atherosclerosis and cancer. J Physiol 2006;574:63–71

47. Dehnavi S, Kiani A, Sadeghi M, Biregani AF, Banach M, Atkin SL, et al. Targeting AMPK by Statins: A Potential Therapeutic Approach. Drugs 2021;81:923–33

48. Ishida CT, Zhang Y, Bianchetti E, Shu C, Nguyen TTT, Kleiner G, et al. Metabolic Reprogramming by Dual AKT/ERK Inhibition through Imipridones Elicits Unique Vulnerabilities in Glioblastoma. Clin Cancer Res 2018;24:5392–406

49. Shen P, Reineke LC, Knutsen E, Chen M, Pichler M, Ling H, et al. Metformin blocks MYC protein synthesis in colorectal cancer via mTOR-4EBP-eIF4E and MNK1-eIF4G-eIF4E signaling. Mol Oncol 2018;12:1856–70

50. Zhang X, Mofers A, Hydbring P, Olofsson MH, Guo J, Linder S, et al. MYC is downregulated by a mitochondrial checkpoint mechanism. Oncotarget 2017;8:90225–37

51. Phang JM, Liu W, Hancock C, Christian KJ. The proline regulatory axis and cancer. Front Oncol 2012;2:60

52. Craze ML, Cheung H, Jewa N, Coimbra NDM, Soria D, El-Ansari R, et al. MYC regulation of glutamine-proline regulatory axis is key in luminal B breast cancer. Br J Cancer 2018;118:258–65

53. Morrish F, Hockenbery D. MYC and mitochondrial biogenesis. Cold Spring Harb Perspect Med 2014;4

54. Iommarini L, Porcelli AM, Gasparre G, Kurelac I. Non-Canonical Mechanisms Regulating Hypoxia-Inducible Factor 1 Alpha in Cancer. Front Oncol 2017;7:286

55. Wang JC, Li XX, Sun X, Li GY, Sun JL, Ye YP, et al. Activation of AMPK by simvastatin inhibited breast tumor angiogenesis via impeding HIF-1α-induced pro-angiogenic factor. Cancer Sci 2018;109:1627–37

56. Das B, Pal B, Bhuyan R, Li H, Sarma A, Gayan S, et al. Regulates the. Cancer Res 2019;79:4015–25

57. Jaggupilli A, Ly S, Nguyen K, Anand V, Yuan B, El-Dana F, et al. Metabolic stress induces GD2. Br J Cancer 2021

58. Ding J, Kuo ML, Su L, Xue L, Luh F, Zhang H, et al. Human mitochondrial pyrroline-5-carboxylate reductase 1 promotes invasiveness and impacts survival in breast cancers. Carcinogenesis 2017;38:519–31

59. Geng P, Qin W, Xu G. Proline metabolism in cancer. Amino Acids 2021;53:1769–77

60. D’Aniello C, Patriarca EJ, Phang JM, Minchiotti G. Proline Metabolism in Tumor Growth and Metastatic Progression. Front Oncol 2020;10:776

61. You D, Du D, Zhao X, Li X, Ying M, Hu X. Mitochondrial malic enzyme 2 promotes breast cancer metastasis via stabilizing HIF-1α under hypoxia. Chin J Cancer Res 2021;33:308–22

62. Nishimura T, Nakata A, Chen X, Nishi K, Meguro-Horike M, Sasaki S, et al. Cancer stem-like properties and gefitinib resistance are dependent on purine synthetic metabolism mediated by the mitochondrial enzyme MTHFD2. Oncogene 2019;38:2464–81

63. Bernhardt S, Bayerlová M, Vetter M, Wachter A, Mitra D, Hanf V, et al. Proteomic profiling of breast cancer metabolism identifies SHMT2 and ASCT2 as prognostic factors. Breast Cancer Res 2017;19:112

64. Siddiqui A, Gollavilli PN, Schwab A, Vazakidou ME, Ersan PG, Ramakrishnan M, et al. Thymidylate synthase maintains the de-differentiated state of triple negative breast cancers. Cell Death Differ 2019;26:2223–36

65. Trachootham D, Alexandre J, Huang P. Targeting cancer cells by ROS-mediated mechanisms: a radical therapeutic approach? Nat Rev Drug Discov 2009;8:579–91

66. Rather GM, Pramono AA, Szekely Z, Bertino JR, Tedeschi PM. In cancer, all roads lead to NADPH. Pharmacol Ther 2021;226:107864

67. Praharaj PP, Panigrahi DP, Bhol CS, Patra S, Mishra SR, Mahapatra KK, et al. Mitochondrial rewiring through mitophagy and mitochondrial biogenesis in cancer stem cells: A potential target for anti-CSC cancer therapy. Cancer Lett 2021;498:217–28

68. El Ansari R, McIntyre A, Craze ML, Ellis IO, Rakha EA, Green AR. Altered glutamine metabolism in breast cancer; subtype dependencies and alternative adaptations. Histopathology 2018;72:183–90

69. Timmerman LA, Holton T, Yuneva M, Louie RJ, Padró M, Daemen A, et al. Glutamine sensitivity analysis identifies the xCT antiporter as a common triple-negative breast tumor therapeutic target. Cancer Cell 2013;24:450–65

70. Gross MI, Demo SD, Dennison JB, Chen L, Chernov-Rogan T, Goyal B, et al. Antitumor activity of the glutaminase inhibitor CB-839 in triple-negative breast cancer. Mol Cancer Ther 2014;13:890–901

71. Prabhu VV, Morrow S, Rahman Kawakibi A, Zhou L, Ralff M, Ray J, et al. ONC201 and imipridones: Anti-cancer compounds with clinical efficacy. Neoplasia 2020;22:725–44

72. Luo B, Ma Y, Zhou Y, Zhang N, Luo Y. Human ClpP protease, a promising therapy target for diseases of mitochondrial dysfunction. Drug Discov Today 2021;26:968–81

73. Wagner J, Kline CL, Ralff MD, Lev A, Lulla A, Zhou L, et al. Preclinical evaluation of the imipridone family, analogs of clinical stage anti-cancer small molecule ONC201, reveals potent anti-cancer effects of ONC212. Cell Cycle 2017;16:1790–9

