## Supplemental materials for "Mitochondrial Matrix Protease ClpP Agonists inhibit Cancer Stem Cell Function in Breast Cancer Cells by Disrupting Mitochondrial Homeostasis"

Greer *et al.*

**Supplementary Methods**

**Supplementary Figures (Fig.S1-S13)**

**Supplementary Table S1**

#### Supplementary Methods

##### *RNA extraction from 15 breast cancer cell lines for RNAseq*

Breast cancer cell lines used were ER+ (ZR75-1, HCC1500, MCF7, T47D), HER2 amplified (BT474, AU565, HCC1954, MB453), TNBC (BT20, HCC1937, MB468, HCC38, MB436, Hs578T, MB231). Original sources of each cell lines and validation of authenticity of cell lines are shown in previous report when RNA was harvested (1).Each breast cancer cell line was grown in 10 cm cell culture plate, trypsinized and centrifuged at 310 x g for 5 minutes, then transferred to 1.5 ml tube, washed with PBS twice, snap frozen with ethanol-dry ice bath. RNA was isolated from cells using TRIZOL reagent as recommended by the manufacturer.

##### *Library preparation and Illumina sequencing for RNAseq*

1 µg RNA per sample was used as the input material for the RNA-seq. RIN (RNA integrity number) values of RNA samples were evaluated using an Agilent 2200 TapeStation system (Agilent Technologies, Santa Clara, CA, USA). Sequencing libraries were generated using NEBNext® rRNA Depletion Kit and NEBNext® Ultra™ Directional RNA Library Prep Kit for Illumina (NEB, USA) following the manufacturer's instructions. The libraries were sequenced on an Illumina HiSeq 2000 platform.

**References**

1. Greer YE, Gilbert SF, Gril B, Narwal R, Peacock Brooks DL, Tice DA, *et al.* MEDI3039, a novel highly potent tumor necrosis factor (TNF)-related apoptosis-inducing ligand (TRAIL) receptor 2 agonist, causes regression of orthotopic tumors and inhibits outgrowth of metastatic triple-negative breast cancer. Breast Cancer Res **2019**;21:27

### Fig.S1

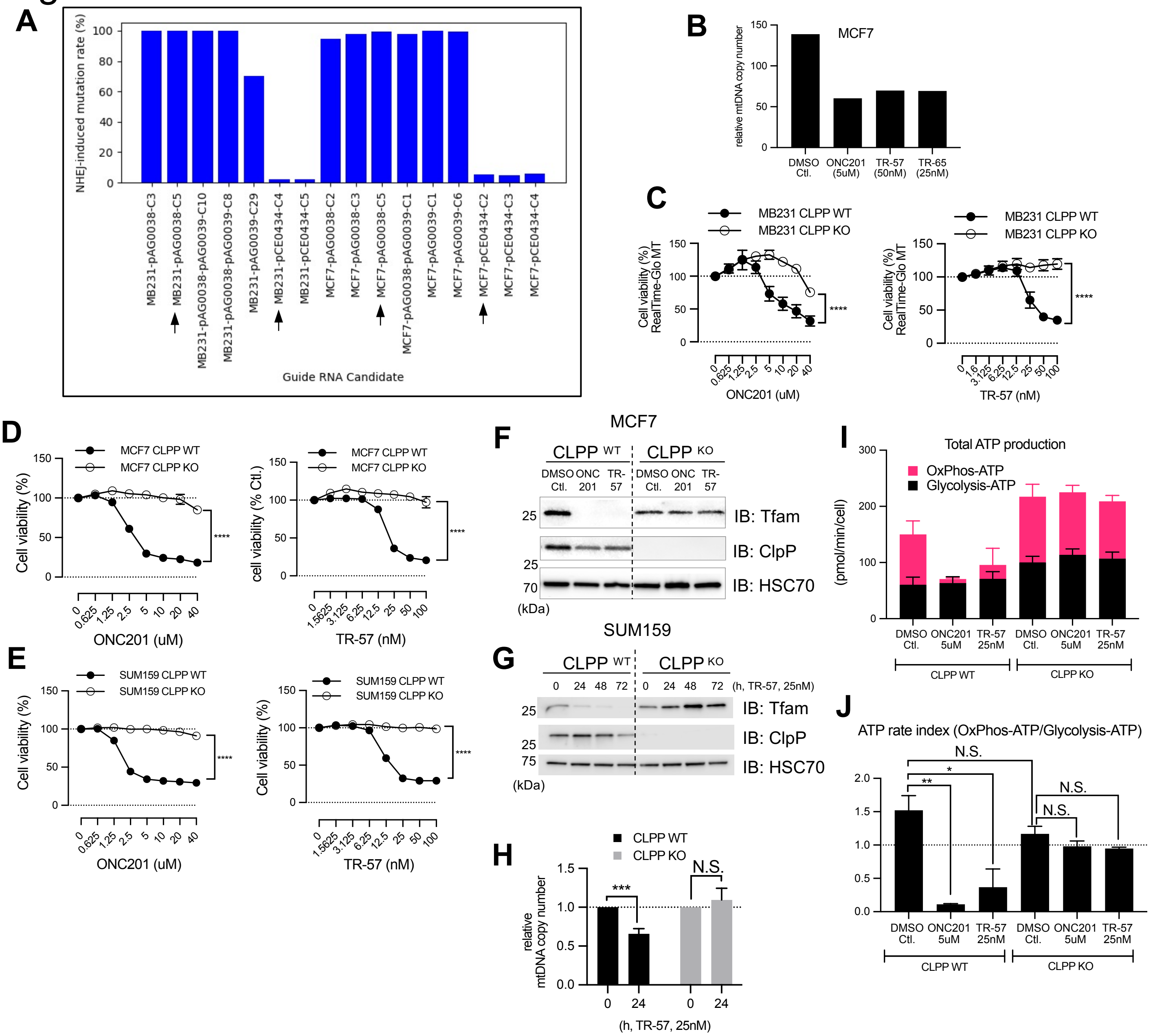


**Fig.S1 ClpP agonists inhibit cell viability and OxPhos in breast cancer cells in a CLPP-dependent manner.**
**A.** The mutation rates of CLPP KO in MB231 and MCF7 cell lines generated by CRISPR-Cas9 system. The cell lines used for the study with arrows. MB231-pCE0434-C4 was used as CLPP WT, MB231-pAG0038-C5 was used as CLPP KO. MCF7-pCE0434-C2 was used as CLPP KO and MCF7-pAG0038-C5 was used as CLPP KO. **B**. Relative mtDNA copy number in MCF7 treated with ClpP agonists for 48h. **C**. RealTime-Glo MT assay with MB231 CLPP WT and KO cell lines treated with ONC201 or TR-57 for 72h. Data shown as ave +/- SEM of 3 independent experiments. ****$p$<0.0001, 2-way ANOVA**. D**. CellTiter-Glo 2.0 assay with MCF7 CLPP WT and KO cells. Data shown as ave +/- SEM of 3 independent experiments. ****$p$<0.0001, 2-way ANOVA. **E.** CellTiter-Glo 2.0 assay with SUM159 CLPP WT and KO cells. Data shown as ave +/- SEM of 3 independent experiments. ****$p$<0.0001. **F**. Western blotting of MCF7 CLPP WT and KO cells treated with ONC201 (5uM) or TR-57 (25nM) for 72h. **G**. Western blotting of SUM159 CLPP WT and KO cells treated with TR-57 (25nM) for 72h. **H**. mtDNA copy numbers in MB231 CLPP WT and KO treated with TR-57 for 24h. Data shown as ave +/- SD of 3 independent experiments. ***$p$<0.001, N.S.; not significant. Student's *t* test. **I.** Seahorse ATP rate assay of SUM159 CLPP WT and KO cell lines treated with DMSO Ctl, ONC201, or TR-57 for 24h. Data shown as ave+/-SEM. *$p$<0.05, **$p$<0.01, N.S.; not significant, Student's *t*-test. **J.** ATP rate index obtained with ATP rate assay shown in Fig.S1I.

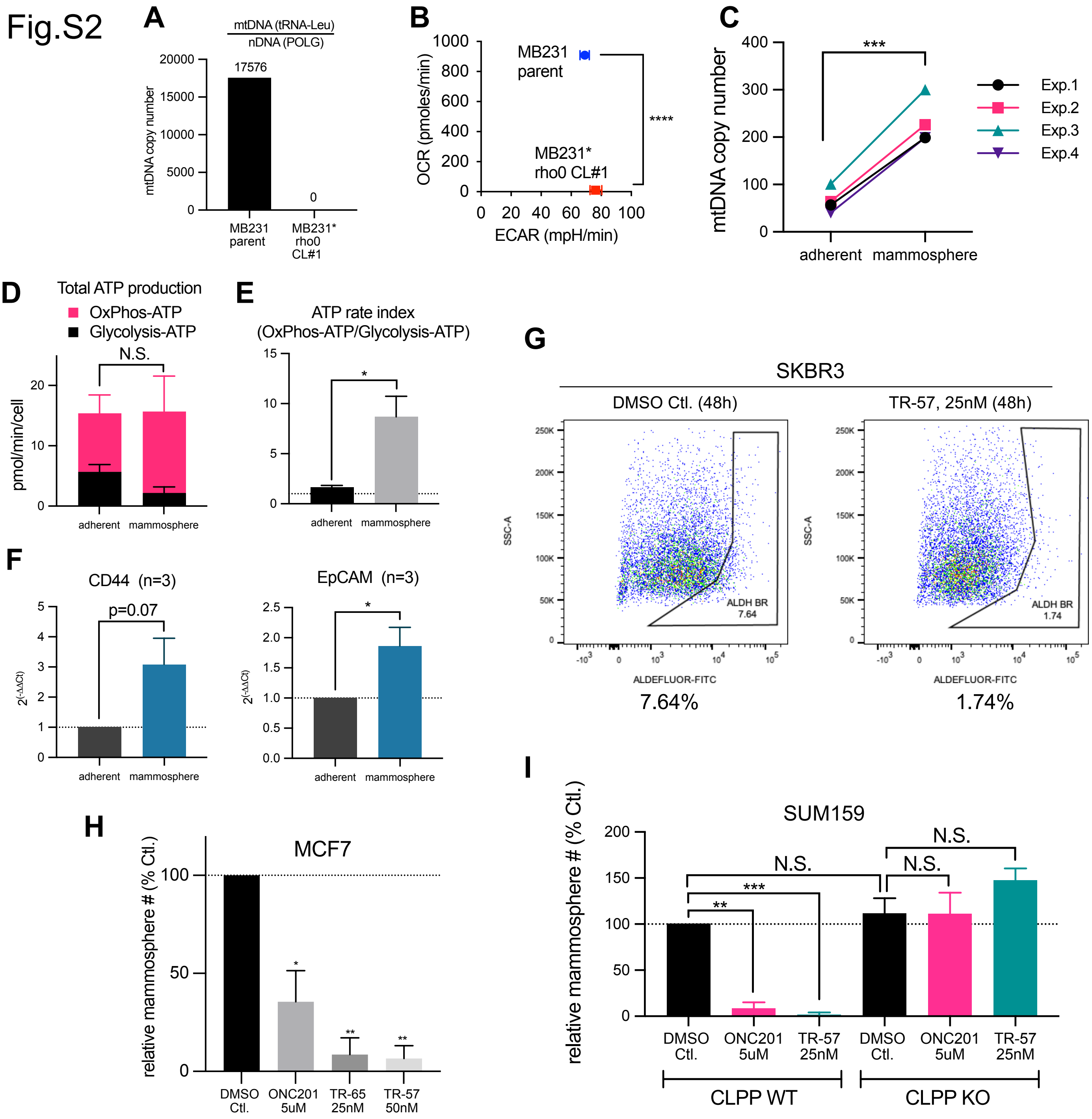


**Fig.S2 Mitochondria are critical for mammosphere formation and ClpP agonists inhibit CSC function *in vitro*.**
**A.** mtDNA copy number qPCR of MB231 parental cells and rho0 cells. **B.** OCR/ECAR profiling of MB231 parental cells and rho0 cells, analyzed with XF analyzer. ****$p$<0.0001, Student's $t$-test. **C.** mtDNA copy number compared between MCF7 adherent cells and mammosphere. ***$p$<0.001, Paired $t$-test. **D&E.** XF analyzer ATP rate assays comparing MCF7 parental cells and mammosphere. Data shown as ave+/- SEM, summary of 3 independent experiments. *$p$<0.05, N.S.; not significant. Student's $t$-test. **F.** qPCR of stem cell markers in MCF7 adherent cells and mammosphere. Data shown as ave+/-SEM, summary of 3 independent experiments. *$p$<0.05, Student's $t$-test. **G.** ALDEFLUOR assays of SKBR3 cells treated with DMSO Ctl. or TR-57 for 48h. Numbers shown below indicate ALDH Bright cells considered as CSC population. **H.** Mammosphere formation assays of MCF7 cells treated with ClpP agonists. Data shown as ave+/-SEM, summary of 2 independent experiments. $p$=0.006, One-way ANOVA, *$p$<0.05, **$p$<0.01, Dunnett's multiple comparisons test (compared with Ctl.). **I.** Mammosphere formation assays performed with SUM159 CLPP WT vs KO cell lines. Data shown as ave+/-SD, summary of 2 independent experiments. **$p$<0.01, ***$p$<0.001, N.S.; not significant. Student's $t$-test.

Fig.S3

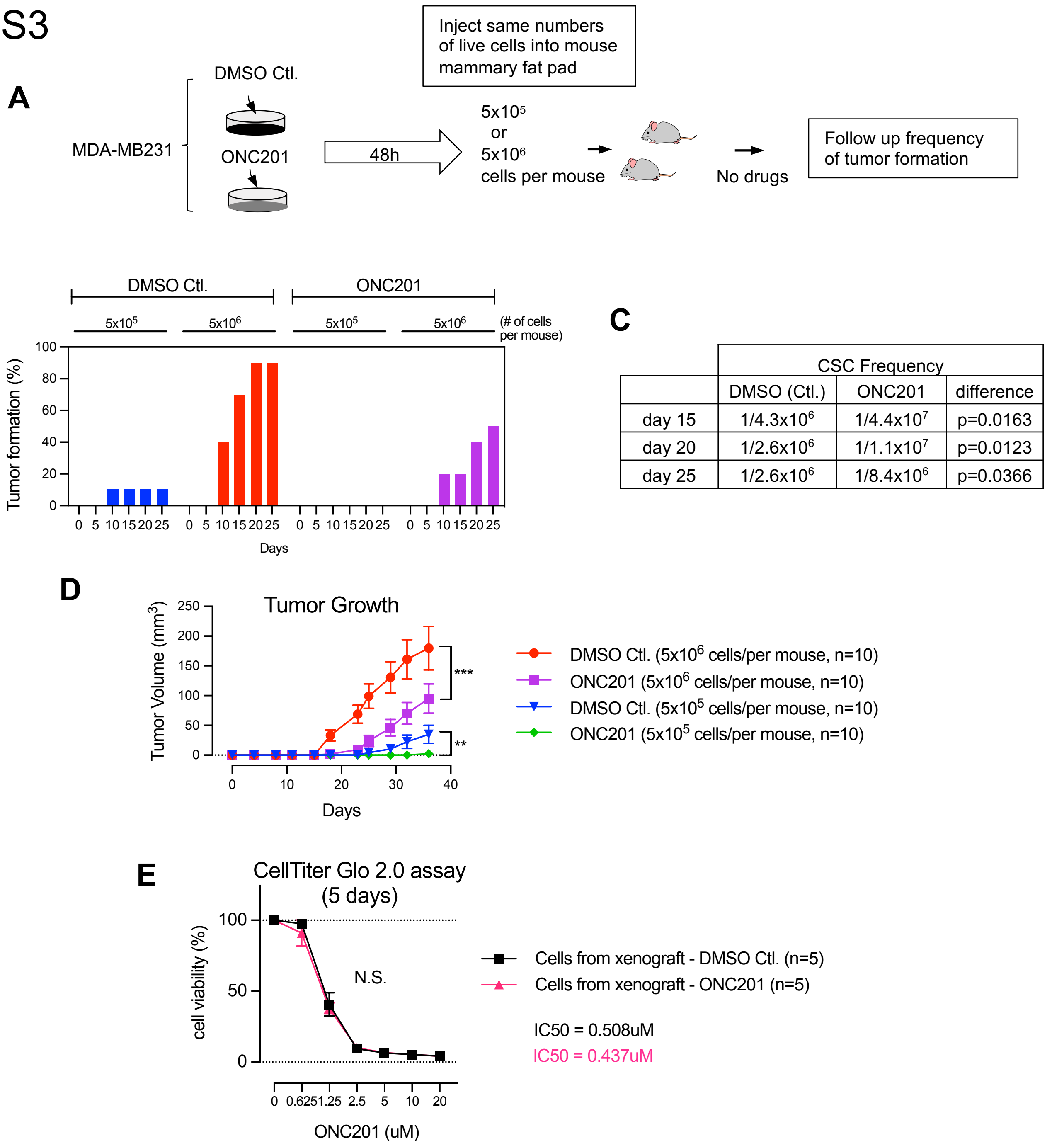


C

| | CSC Frequency | | |
|---|---|---|---|
| | DMSO (Ctl.) | ONC201 | difference |
| day 15 | 1/4.3x10$^6$ | 1/4.4x10$^7$ | p=0.0163 |
| day 20 | 1/2.6x10$^6$ | 1/1.1x10$^7$ | p=0.0123 |
| day 25 | 1/2.6x10$^6$ | 1/8.4x10$^6$ | p=0.0366 |

**Fig.S3 ClpP agonists inhibit tumor initiation *in vivo.***
A. Experimental procedure illustrating the 1$^{st}$ *in vivo* tumorigenicity experiment. **B**. Tumor formation (%) in each group at different time points. Day 0 is the day of cell injection to mammary fat pad. **C**. CSC frequency between control and ONC201-treated groups at different time points was determined using ELDA software. **D.** Tumor growth curve in the 1st experiment up to Day 36. Data shown as ave+/-SEM of tumor size from each group. **$p$<0.01, ***$p$<0.001, 2-way ANOVA. **E.** CellTiter-Glo 2.0 assays. Five tumors grown in 5x10$^6$ cells/mouse (DMSO Ctl. group) and 5 tumors grown 5x10$^6$ cells/mouse (ONC201-group) from the 1$^{st}$ tumorigenicity experiment were collected and human cells were isolated. The data is shown as ave+/-SD. N.S.; not significant, 2-way ANOVA.

Fig.S4

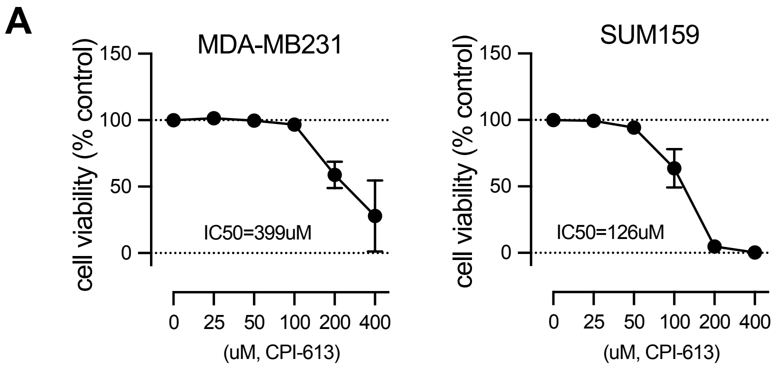


**Fig.S4 The TCA cycle inhibitor CPI-613 downregulates cell viability in MB231 and SUM159 cell lines.**
**A.** CellTiter-Glo 2.0 assays of MB231 (left) and SUM159 (right) cells treated with CPI-613 for 3 days. Data shown as ave+/-SEM, summary of multiple independent experiments.

Fig.S5

A

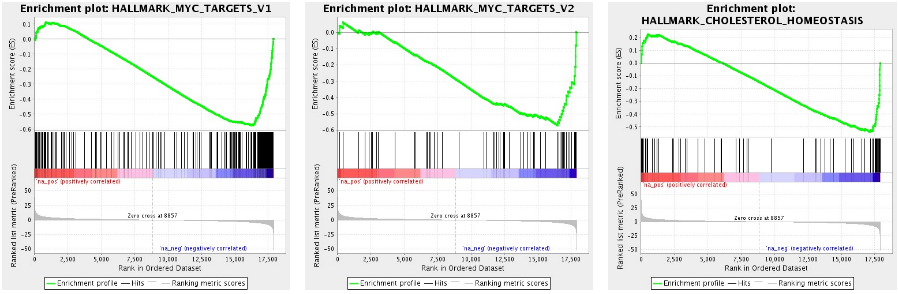

| Geneset name | ES | NES | NOM p-val | FDR q-value |
|---|---|---|---|---|
| HALLMARK_MYC_TARGETS_V1 | -0.5755139 | -2.2348235 | 0 | 0 |
| HALLMARK_MYC_TARGETS_V2 | -0.5718771 | -1.8871752 | 0 | 2.44E-04 |
| HALLMARK_CHOLESTEROL_HOMEOSTASIS | -0.5405826 | -1.8157017 | 0 | 3.55E-04 |

B

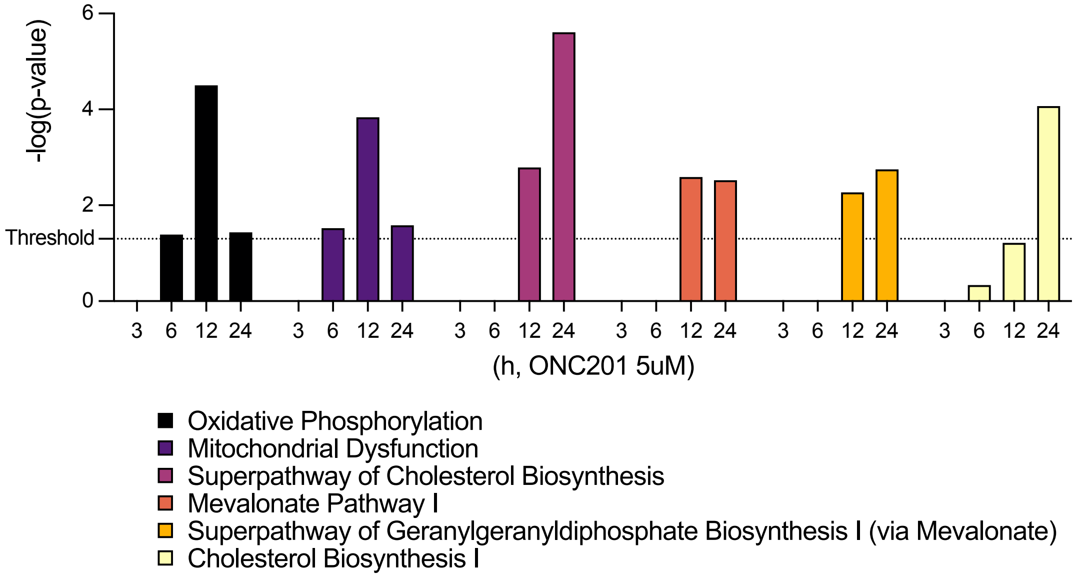


C

| Maps | min(pValue) | Min FDR |
|---|---|---|
| SCAP/SREBP Transcriptional Control of Cholesterol and FA Biosynthesis | 1.176E-14 | 6.140E-12 |
| Transcription_HIF-1 targets | 4.753E-07 | 3.601E-05 |
| Development_YAP/TAZ-mediated co-regulation of transcription | 6.501E-07 | 4.434E-05 |

**Fig.S5 RNAseq indicates that ONC201 dysregulates multiple signaling pathways and proteins critical for CSC maintenance.**
**A.** GSEA plots indicating hallmark genesets negatively regulated by ONC201 24h treatment in MB231 cells. The data suggests transcripts involved with Myc target genes and cholesterol pathway. The table shows 4 key statistics for the GSEA. ES: enrichment score, NES: Normalized enrichment score, NOM p value: nominal p value, FDR q-value: False discovery rate.
**B.** Ingenuity pathway analysis (IPA) indicating that OxPhos and cholesterol synthesis pathways are dysregulated by ONC201.
**C.** Metacore enrichment analysis indicating that ONC201 24h treatment significantly dysregulates cholesterol and fatty acid synthesis pathway, transcription of HIF1 targets, and YAP/TAZ pathway.

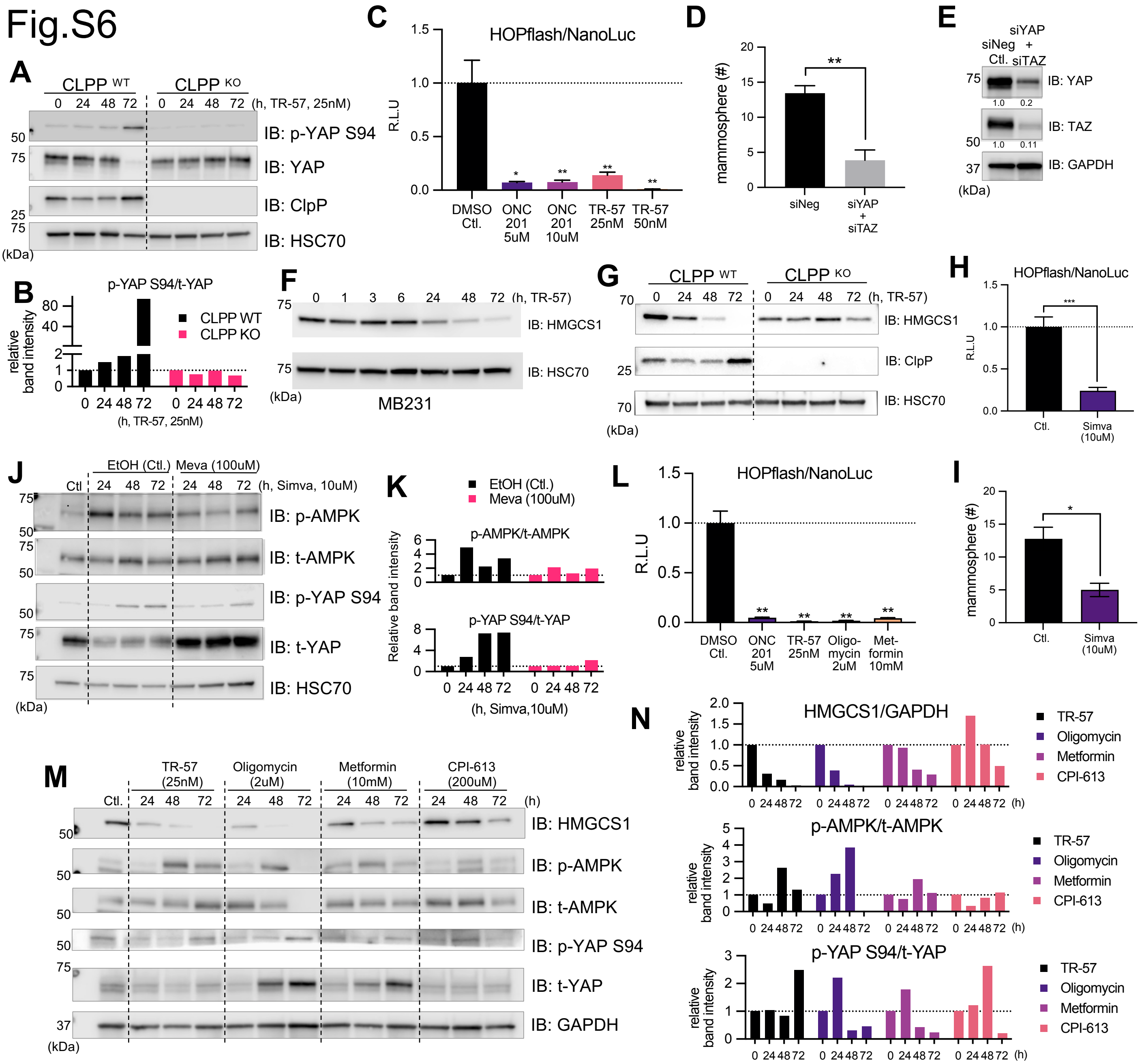


**Fig.S6 ClpP agonists and other mitochondria-targeting drugs inhibit mevalonate pathway via AMPK-dependent down-regulation of YAP/TAZ pathway.**
**A.** Representative immunoblot showing that TR-57 induces YAP phosphorylation Ser94 in a CLPP-dependent manner in MB231 cells. **B.** Relative band intensities of phospho-YAP (Ser94) normalized with total YAP from data presented in panel A. **C.** HOPflash reporter assay with SUM159 cells after treated with ClpP agonists for 72h. Data shown as ave+/-SD. **D.** Mammosphere formation assays with MB231 cells transfected with negative ctl. or YAP/TAZ siRNA. Data shown as ave+/-SEM, summary of 3 independent experiments. **E.** Representative immunoblot data for panel D. **F.** Representative Western blot showing time-dependent effect of TR-57 (25nM) on HMGCS1 in MB231 cells. **G**. Representative Western blot showing time-dependent effect of TR-57 (25nM) on HMGCS1 in MB231 CLPP WT and KO cells. **H.** Representative HOPflash reporter assay with SUM159 cells after treated with simvastatin for 48h. **I.** Mammosphere formation assays with MB231 cells treated with simvastatin. Drugs were added every 2-3 days. Data shown as ave+/-SEM, summary of 3 independent experiments. **J.** Representative immunoblot showing that simvastatin induces AMPK activation and phosphorylation of YAP at Ser94, and it was inhibited by reversed with mevalonolactone in MB231 cells. **K.** Relative band intensities of phospho-AMPK normalized with total AMPK, and phospho-YAP (Ser94) normalized with total YAP shown in panel J. **L**. Representative HOPflash reporter assay with SUM159 after treated with multiple mitochondria-targeting drugs for 72h. Data shown as ave+/-SD. **M.** Representative immunoblot showing the effect of multiple mitochondria-targeting drugs on HMGCS1, AMPK and YAP in MB231 cells. **N.** Relative band intensities of HMGCS1 normalized with GADPH, phospho-AMPK normalized with total AMPK, and phospho-YAP (Ser94) normalized with total YAP shown in panel **M**.

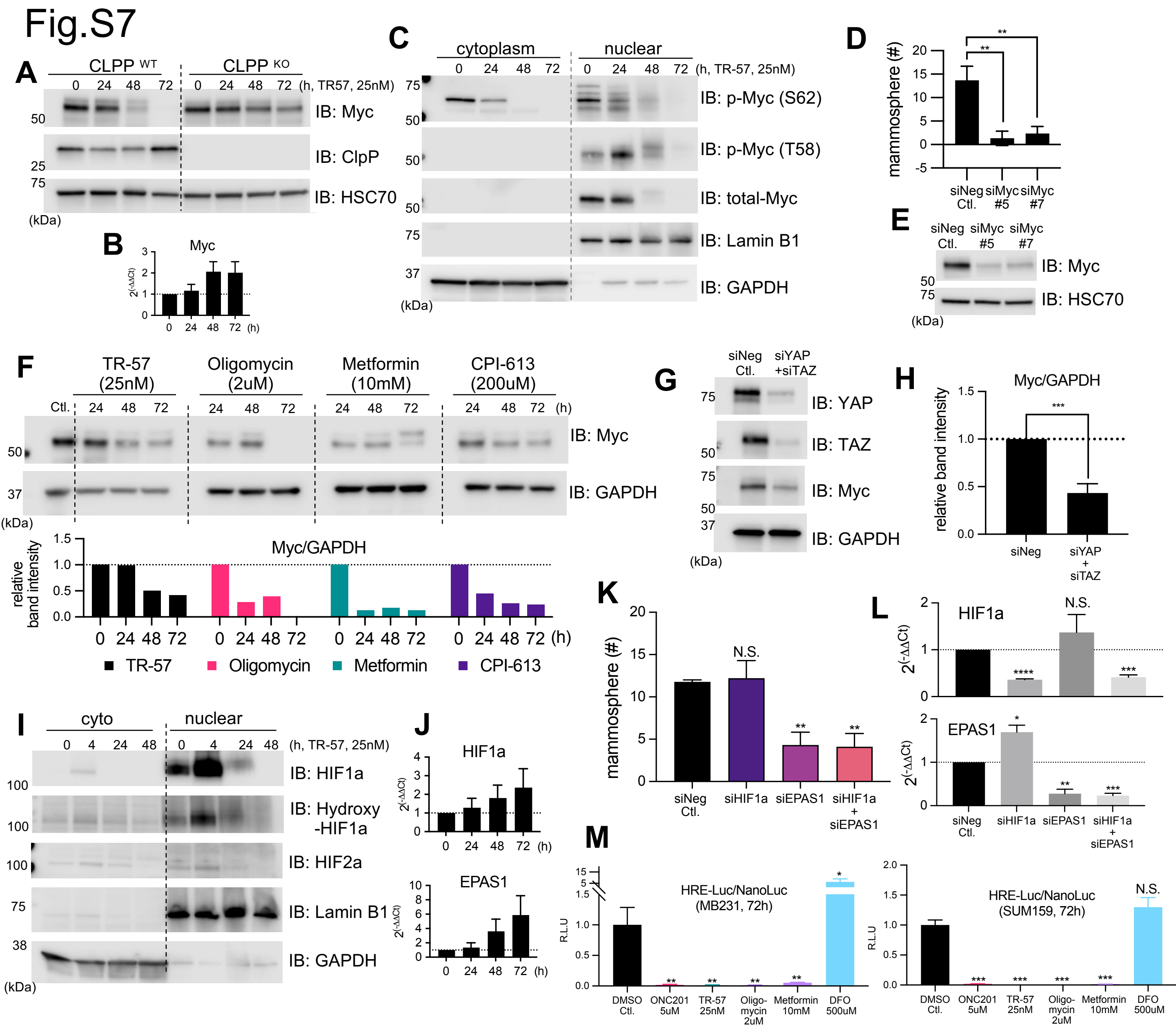


**Fig.S7 ClpP agonist and other mitochondria targeting drugs downregulate Myc and HIF expression.**
**A.** Representative immunoblot showing TR-57 downregulates Myc in MB231 cells in a CLPP-dependent manner. **B.** Time-dependent effect of TR-57 on Myc transcript in MB231 cells, analyzed by qPCR. Data shown as ave+/-SEM, summary of 4 independent experiments. Not significant, but there was a trend of increase over the time. **C.** Representative immunoblot showing that TR-57 promoted Myc phosphorylation at Thr58 and downregulated total Myc level in nuclear fraction in MB231 cells. **D.** The effect of knockdown of Myc on mammosphere formation assays in MB231 cells. One of two independent experiments using two different siRNA against Myc. **E.** Accompanied immunoblot data for panel D showing Myc knockdown 48h after siRNA transfection. **F.** Representative immunoblot showing multiple mitochondria-targeting drugs downregulate Myc. Relative band intensities of Myc normalized with GADPH are shown in the lower panel. **G.** The effect of YAP/TAZ knockdown on Myc expression in MB231 cells. Cells were harvested 48h after siRNA transfection. One of 3 independent experiments. **H.** Relative band intensities of Myc normalized with GAPDH. Data shown as ave+/- SD, summary of 3 independent experiments. ****$p$<0.0001, Student's $t$-test. **I.** Time-dependent effect of TR-57 on HIF1α and HIF2α in SUM159 cells. GAPDH and Lamin B1 were used as cytosolic and nuclear fraction markers, respectively. **J.** Time-dependent effect of TR-57 on HIF1α and EPAS1 transcripts detected by qPCR. MB231 cells. Data shown as ave+/-SEM, summary of 4 independent experiments. Not significant, but there was a trend of increasing level of the transcripts. **K.** Mammosphere formation assay with MB231 cells transfected with HIF1α/EPAS1 siRNA. Data shown as ave+/-SEM, summary of 3 independent experiments. **$p$<0.01, N.S.; not significant, Student's $t$-test. **L.** Accompanied qPCR data for panel K, showing the effect of HIF1a/EPAS1 siRNA, Data shown as ave+/-SEM, summary of 3 independent experiments. Cells were collected 8 days after siRNA transfection. **M**. ClpP agonists, as well as other mitochondria-targeting drugs inhibit HRE-Luc in both MB231 and SUM159 cells, 72h treatment. Deferoxamine (DFO), an iron-chelator, was used as a positive control. *$p$<0.05, **$p$<0.01, ***$p$<0.001, N.S.; not significant, Student's $t$-test.

Fig.S8

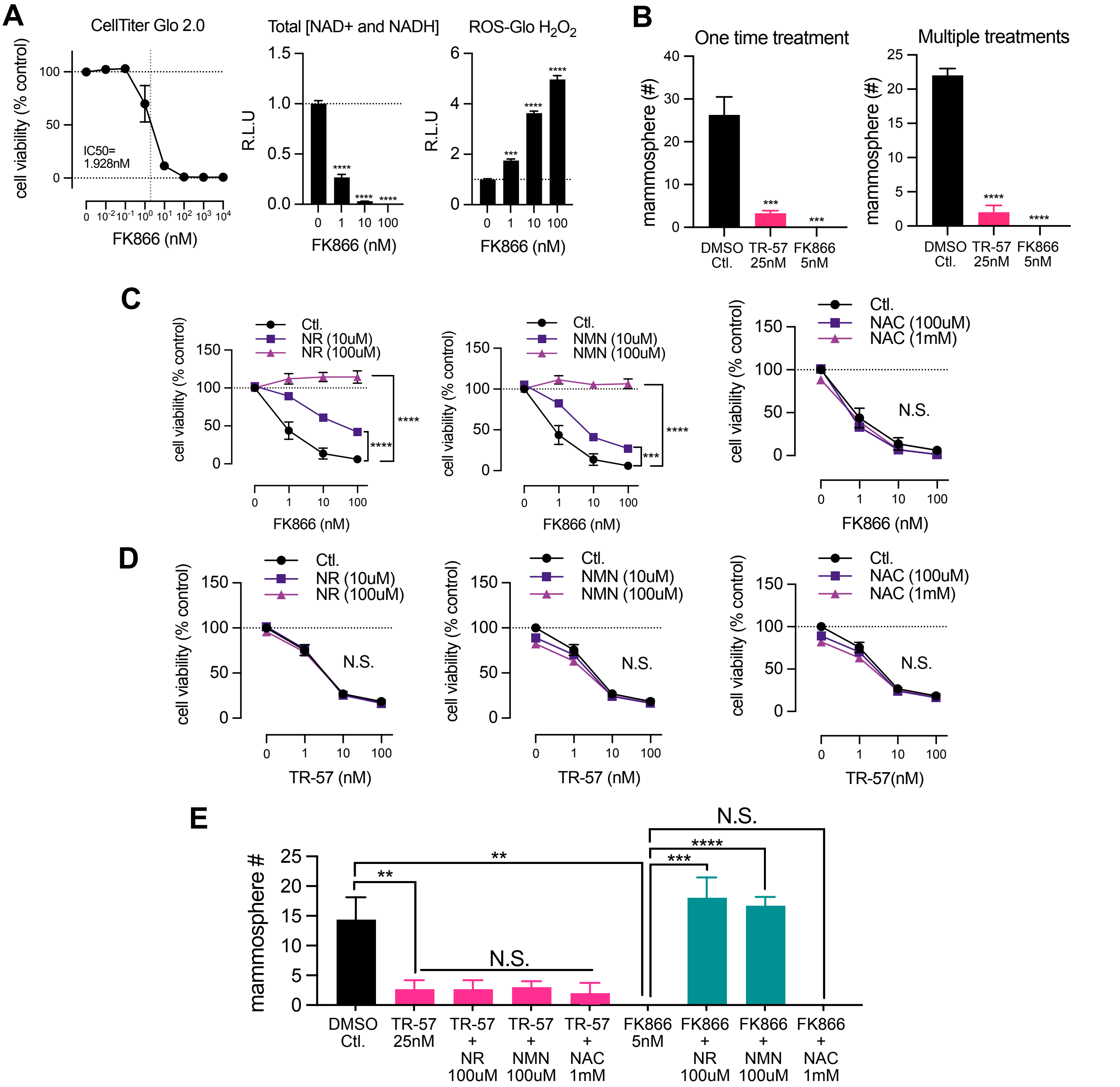


**Fig.S8 ClpP agonist downregulates NAD(P)/NAD(P)H and induces oxidative stress**

**A.** CellTiter-Glo 2.0, total NAD+ and NADH, and ROS-Glo $H_2O_2$ assays of MB231 cells treated with FK866 for 5 days. Data shown as ave+/-SEM. ***$p$<0.001, ****$p$<0.0001, N.S.; not significant. Student's $t$-test. **B.** Mammosphere formation assay with MB231 cells treated with TR-57 or FK866. Data shown as ave+/-SD. ***$p$<0.001, ****$p$<0.0001, N.S.; not significant. Student's $t$-test. **C.** CellTiter-Glo2.0 assays of MB231 cells treated with FK866 with or without NR, NMN, NAC for 5 days. Data shown as ave+/-SD. ***$p$<0.001, ****$p$<0.0001, N.S.; not significant. 2-way ANOVA. **D.** CellTiter-Glo2.0 assays of MB231 cells treated with TR-57 with or without NR, NMN, NAC for 5 days. Data shown as ave+/-SD. N.S.; not significant. 2-way ANOVA. **E.** Mammosphere formation assays of MB231 cells treated with TR-57 or FK866 with or without NR, NMN, NAC. **$p$<0.001, ***$p$<0.001, ****$p$<0.0001, N.S., not significant. Student's $t$-test.

Fig.S9

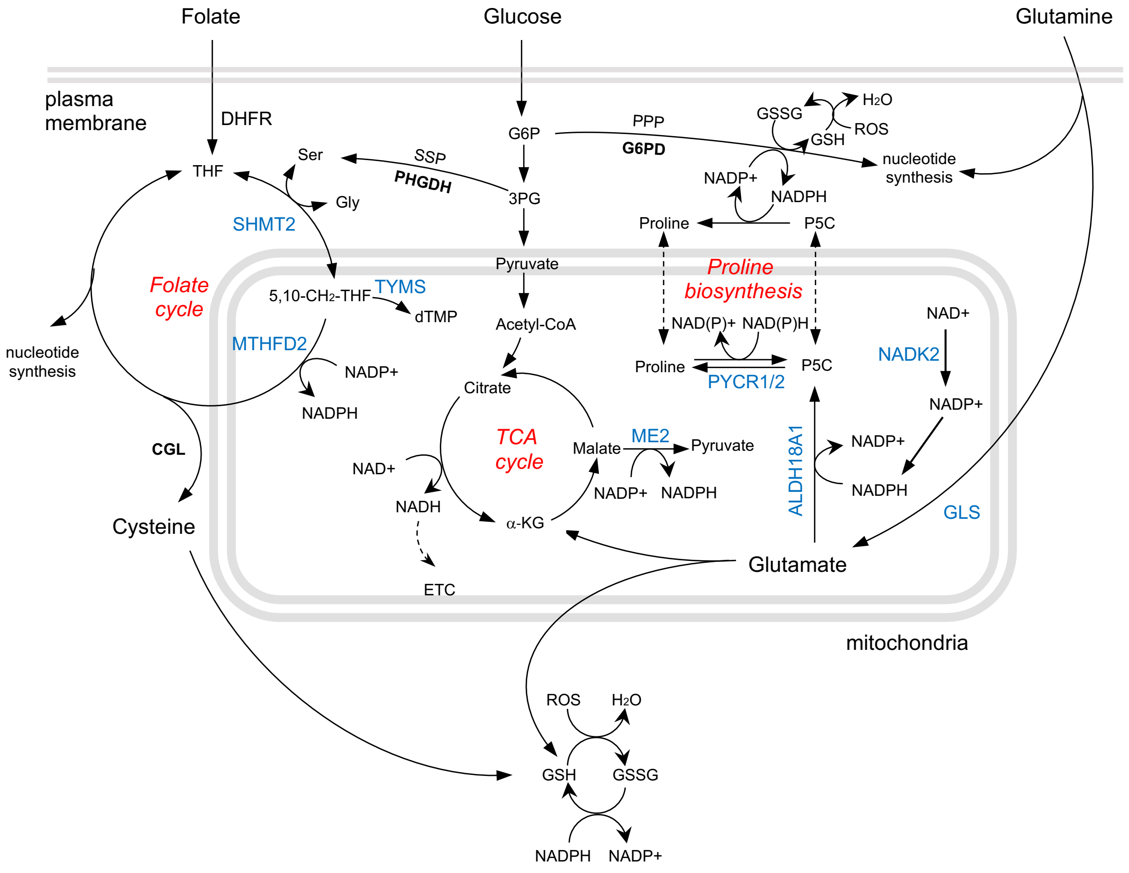


**Fig.S9 ClpP agonists downregulated multiple enzymes involved with glutamine-proline axis and NADPH-generation in mitochondria.**

The enzymes downregulated by ClpP agonists are highlighted with blue. Metabolic pathways dysregulated by ClpP agonists are highlighted with red.

### Fig.S10

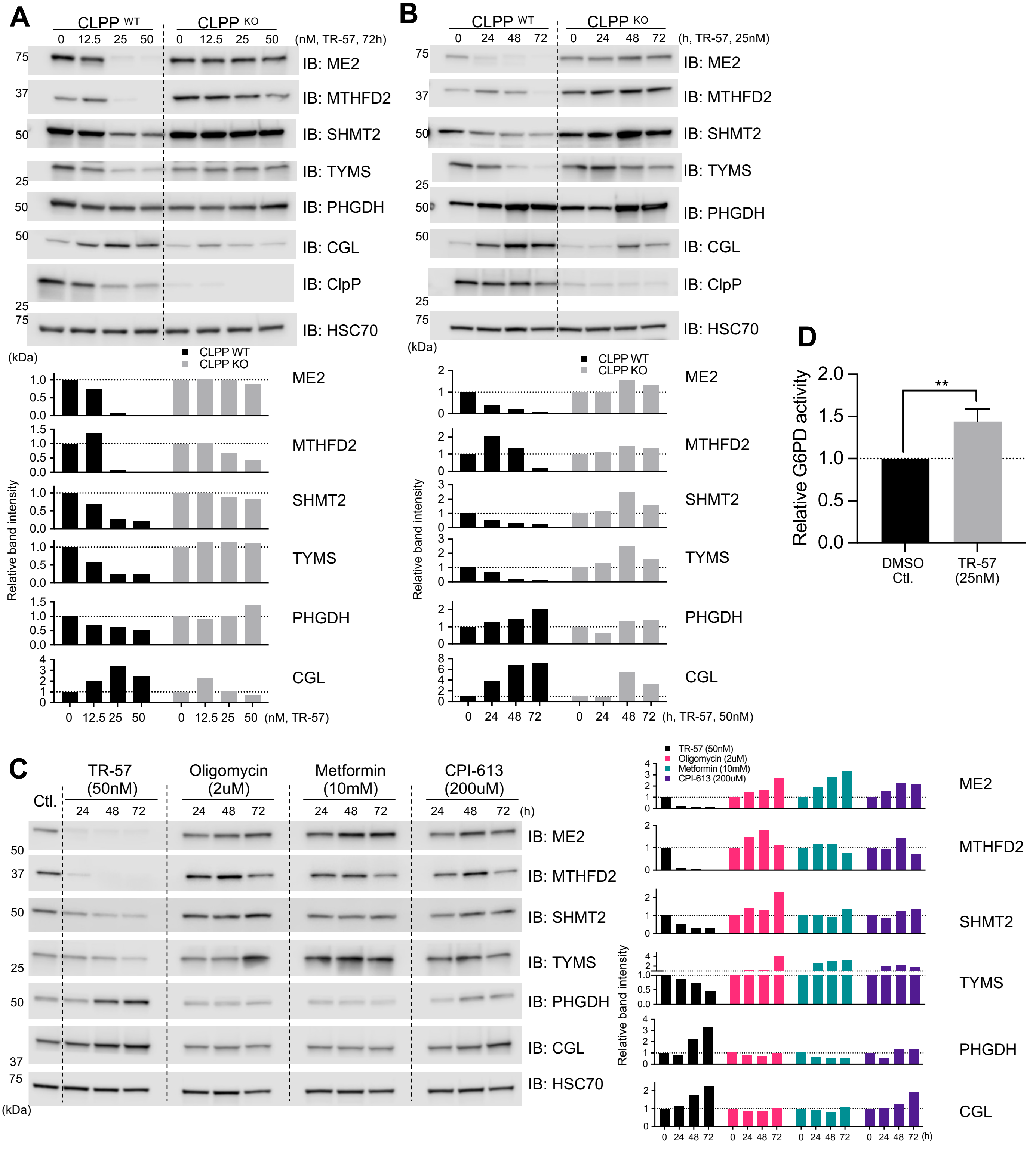


**Fig.S10 ClpP agonist inhibits folate-mediated one carbon metabolism.**
**A&B.** Immunoblots showing dose (**A**) and time (**B**)-dependent effects of TR-57 on enzymes involved with FOCM, serine synthesis pathway in SUM159 CLPP WT and KO cell lines. Representative data from multiple experiments are shown. Relative band intensities of each protein are shown in the lower panels. **C**. Representative immunoblots comparing the time-dependent effect of various mitochondria-targeting drugs on FOCM and serine synthesis pathway in MB231 cells. Relative band intensities of each protein are shown in the right. **D.** G6PD enzymatic activity assays with MB231 cells treated with DMSO Ctl. or TR-57 for 72h. Data shown as ave+/-SD, summary of 3 independent experiments. ***p*<0.01, Student's *t*-test.

Fig.S11

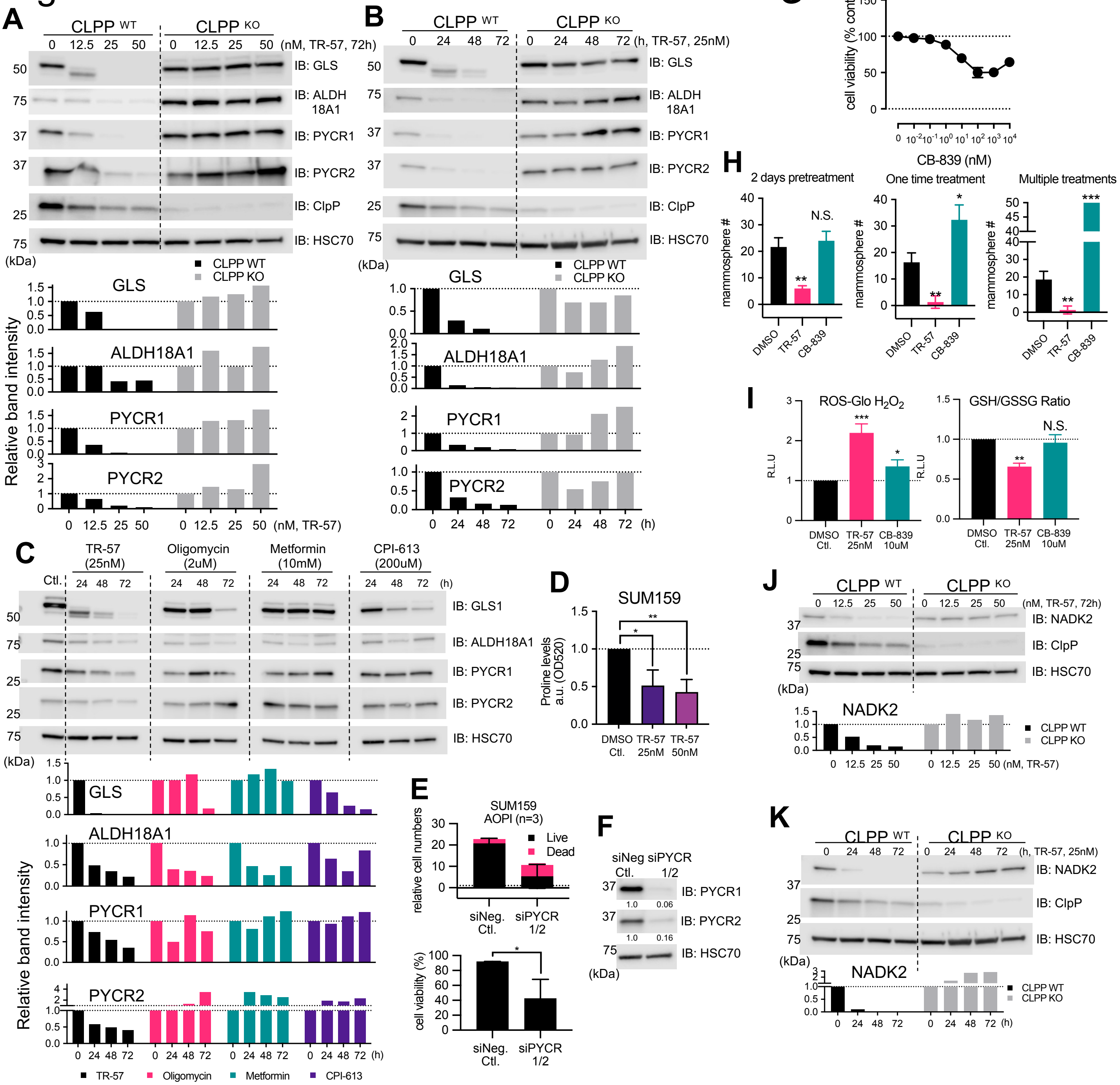


**Fig.S11 ClpP agonist downregulates proline biosynthesis.**

**A&B.** The dose(**A**) and time(**B**)-dependent effect of TR-57 on enzymes involved with glutamine-proline axis in SUM159 CLPP WT and KO cell lines. Relative band intensities of each protein are shown in the lower panels. Representative data from multiple experiments are shown. **C.** The effects of multiple mitochondria-targeting drugs on enzymes involved with glutamine-proline axis in MB231 cells. Representative data from multiple similar results. Relative band intensities of each protein are shown in the lower panels. **D.** Proline assays of SUM159 cells treated with TR-57 for 72h. Data shown as ave+/-SD, summary of 3 independent experiments. **E.** Relative cell viabilities analyzed with AOPI assays. Cell viability of SUM159 was examined after 3 days of siRNA transfection. Data shown as ave+/-SD, 3 independent experiments. Left: numbers of live and dead cells relative to initial cell numbers (dotted line) transfected. Right: % of cell viability (live/total cell numbers). ***p*<0.01, Student's *t*-test. **F.** Representative immunoblot of experiments shown in panel E. **G.** CellTiter-Glo 2.0 assay with MB231 cells treated with CB-839 for 72h. Data shown as ave+/-SEM, summary of 4 independent experiments. **H.** Mammosphere formation assays of MB231 cells treated with DMSO Ctl, TR-57 (25nM), or CB-839 (10uM). Three different mammosphere formation assay procedures were used as shown in Fig.4A-C. Data shown ave+/-SD. **p*<0.05, ***p*<0.01, ****p*<0.001, N.S.; not significant, Student's *t*-test. **I.** ROS-Glo $H_2O_2$ and GSH/GSSG ratio assays in MB231 cells, 5 days drug treatment. Data shown as ave+/-SEM, summary of multiple experiments. **p*<0.05, ***p*<0.01, ****p*<0.001, N.S.; not significant, Student's *t*-test. **J.** Dose-dependent effect of TR-57 on NADK2 in SUM159 cells. Relative band intensities of NADK2 normalized with HSC70 is shown in the lower panel. **K.** Time-dependent effect of TR-57 on NADK2 in SUM159 cells. Relative band intensities of NADK2 normalized with HSC70 is shown in the lower panel.

Fig.S12

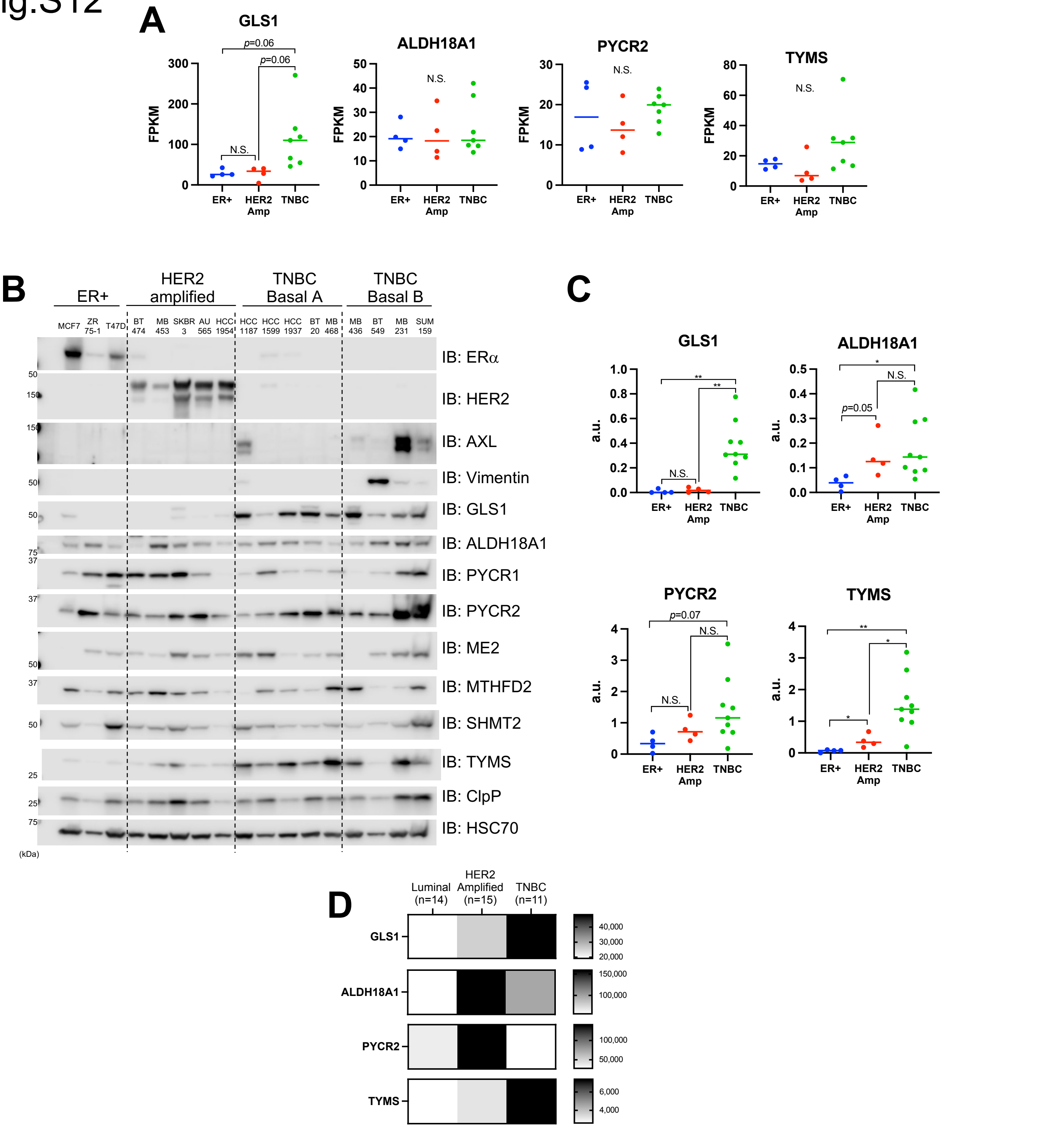


**Fig.S12 The expression of ClpP-targeted enzymes in breast cancer cell lines and breast cancer patients.**
**A**. Comparison of GLS1 Transcript level in 15 breast cancer cell lines representing 3 different molecular subtypes by RNAseq. The subtypes of cell lines used were: ER+ (ZR75-1, HCC1500, MCF7, T47D), HER2 amplified (BT474, AU565, HCC1954, MB453), TNBC (BT20, HCC1937, MB468, HCC38, MB436, Hs578T, MB231). Student's *t*-test. ALDH18A1, PYCR1, PYCR2, TYMS, PYCR1, ME2, MTHFD2, SHMT2, did not show statistical difference. **B&C**. Protein expression of enzymes involved with glutamine-proline axis, ME2 and FOCM across 17 breast cancer cell lines. The subtypes of cell lines used in the Western blot were: ER+ (n=3), HER2 amplified (n=5), TNBC (n=9) *$p<0.05$, **$p<0.01$, N.S.; Not Significant, Student's *t*-test. PYCR1, ME2, MTHFD2, SHMT2, ClpP did not show statistical difference between subtypes, therefore not shown. The bars shown in C indicates the median value. **D.** Comparison of protein expression levels of GLS1, ALDH18A1, PYCR2 and TYMS in breast cancer patients (n: patients numbers) with different molecular subtypes.

Fig.S13

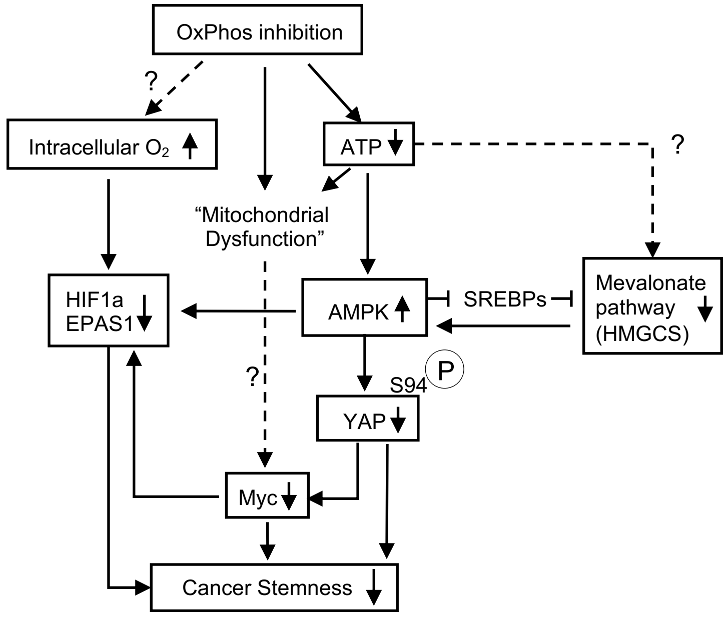


**Fig.S13 OxPhos inhibition dysregulates network of multiple pathways involved with CSC functions.** A diagram illustrating proposed mechanisms how OxPhos inhibition downregulates mevalonate pathway, YAP/TAZ pathway, Myc and HIF pathway, contributing to suppression of CSC functions.

### Supplementary Table S1 (1/3)

| Chemicals, reagents, assay kits | | | |
|---|---|---|---|
| Reagent/Kit | Source | Identifier/catalog number | Stock and storage |
| Cellometer ViaStain™ AOPI Staining Solution | Nexcelcom | CS2-0106-5mL | |
| CellTiter-Glo® 2.0 Cell Viability Assay | Promega | G9242 | |
| Daporinad (FK866, APO866) | SelleckChem | S2799 | 10mM in DMSO, -30$^o$C |
| Deferoxamine mesylate salt (DFO) | Sigma-Aldrich | D9533-1G | |
| Devimistat (CPI-613) | SelleckChem | S2776 | 200mM in DMSO, -30$^o$C |
| Dimethylsulfoxide (DMSO) | Sigma-Aldrich | D2650-100ml | |
| General Proline Assay Kit | MyBiosource | MBS8305379 | |
| GLS1 Inhibitor III, CB-839 | Sigma-Aldrich | 5337170001 | 20mM in DMSO, -30$^o$C |
| Glucose 6 Phosphate Dehydrogenase Assay Kit | Abcam | ab102529 | |
| GSH/GSSG-Glo™ Assay | Promega | V6611 | |
| Hoechst 33342 | ThermoFisher | H3570 | |
| Lipofectamine® 3000 | ThermoFisher Scientific | L3000008 | |
| Lipofectamine® RNAiMax | ThermoFisher Scientific | 13778150 | |
| metformin hydrochloride | Sigma-Aldrich | PHR1084-500mg | 1M in PBS, -30$^o$C |
| mevalonolactone | Sigma-Aldrich | M4667-1G | 100mM in 95% EtOH, -30$^o$C |
| N-acetyl-cysteine | Sigma-Aldrich | A9165 | |
| NAD/NADH-Glo Assay | Promega | G9071 | |
| NADP/NADPH-Glo Assay | Promega | G9081 | |
| Nano-Glo® Dual-Luciferase® Reporter Assay System | Promega | N1610 | |
| Nicotinamide Riboside Chloride (NR, NIAGEN) | SelleckChem | S2935 | |
| oligomycin | Sigma-Aldrich | O4876 | 10mM in DMSO, -30$^o$C |
| ONC201 | Chimerix, Inc (Durham, NC) | https://www.chimerix.com/ | 20mM in DMSO, -30$^o$C |
| puromycin | ThermoFisher | A1113802 | |
| RealTime-Glo™ MT Cell Viability Assay | Promega | G9712 | |
| ROS-Glo™ H2O2 Assay | Promega | G8820 | |
| simvastatin | Sigma-Aldrich | 567020-50MG | 60mM,55% EtOH, 45% 1N NaOH, pH 7.2, -30$^o$C |
| ß-Nicotinamide Mononucleotide (NMN) | SelleckChem | S5259 | |
| TR-57, TR-65 | Madera Therapeutics, LLC (Cary, NC) | http://maderathera.com/ | 50uM in DMSO, -30$^o$C |

| Materials for molecular biology experiments | | |
|---|---|---|
| Reagent/Kit | Source | Identifier/catalog number |
| All Prep DNA/RNA Mini Kit | Qiagen | 80204 |
| DNeasy Blood & Tissue kit | Qiagen | 69504 |
| Vimentin | ThermoFisher Scientific | 11635018 |
| EndoFree Plasmid Maxi Kit (10) | Qiagen | 12362 |
| Power Up SYBR Green Master Mix | ThermoFisher Scientific | A25780 |
| QuantiTect Reverse Transcription Kit | Qiagen | 205313 |
| Quick Ligation Kit | New England Biolabs | M2200S |
| RNeasy Mini Kit (50) | Qiagen | 74104 |
| T4 DNA ligase | New England Biolabs | M0202S |
| T4 Polynucleotide Kinase | New England Biolabs | M0201S |
| TRIZOL | ThermoFisher Scientific | 15596018 |
| Primer for qPCR | Source | Identifier/catalog number |
| CD44 | Qiagen | QT00998333 |
| EPAS1 | Qiagen | QT00069587 |
| EpCAM | Qiagen | QT00000371 |
| GAPDH | Qiagen | QT00079247 |
| HIF1a | Qiagen | QT00083664 |
| Human Mitochondrial DNA (mtDNA) Monitoring Primer Set | Takara Bio USA, Inc | 7246 |
| Myc | Qiagen | QT00035406 |
| ZEB | Qiagen | QT00020972 |
| siRNA | Source | Identifier/catalog number |
| all stars negative control siRNA | Qiagen | 1027281 |
| Hs_EPAS1_5 FlexiTube siRNA | Qiagen | SI02663038 |
| Hs_HIF1A_6 FlexiTube siRNA | Qiagen | SI02664431 |
| Hs_HMGCS1_1 FlexiTube siRNA | Qiagen | SI00033061 |
| HS_MYC_5 Flexitube siRNA | Qiagen | SI00300902 |
| HS_MYC_7 Flexitube siRNA | Qiagen | SI02662611 |
| Hs_PYCR1_12 Flexitube siRNA | Qiagen | SI04983587 |
| Hs_PYCR2_8 FlexiTube siRNA | Qiagen | SI04294745 |
| Hs_WWTR1_1 FlexiTube siRNA (TAZ) | Qiagen | SI00111216 |
| Hs_YAP1_1 FlexiTube siRNA | Qiagen | SI00084546 |
| CRISPR/Cas9 knock out | target sequence | Identifier/catalog number |
| Lenti-CRISPR-V2 backbone | Addgene | 52961 |
| sgRNA1 CLPP target sequence | GCGCCTATGACATCTACTCGCGG | Oligonucleotide: IDT Technologies (Coralville, IA) |
| sgRNA2 CLPP target sequence | GGAGCGCATCGTGTGCGTCATGG | |
| Plasmid DNA | Source | Identifier/catalog number |
| Hop-flash | Addgene | 83467 |
| HRE-Luciferase | Addgene | 26731 |
| pNL1.1.TK[Nluc/TK] | Promega | N1501 |

### Supplementary Table S1 (2/3)

| **Materials for Western blotting** | | |
|---|---|---|
| Primary antibody | Source | Identifier/catalog number |
| ALDH18A1 | Sigma-Aldrich | HPA012604-100UL |
| AMPK | Cell Signaling | 2532 |
| AXL | Santa Cruz Biotechnology | sc-20741 |
| Chameleon Duo Protein Ladder | LICOR | 928-600000 |
| CLPP | Cell Signaling | 14181 |
| Cystathionine gamma-Lyase/CGL | Cell Signaling | 19689S |
| Estrogen Receptor alpha | Cell Signaling | 8644S |
| G6PD | Cell Signaling | 12263S |
| GAPDH | Cell Signaling | 2118 |
| Glutaminase-1/GLS1 | Cell Signaling | 56750 |
| HER2 (c-erb B-2) | NeoMarkers | RB-103-P1 |
| HIF-1 alpha | BD biosciences | 610958 |
| HIF-2α | Novus Biologicals Inc. | NB100-122 |
| HMGCS1 (D5W8F) Rabbit mAb | Cell Signaling | 36877 |
| HSC70 | Santa Cruz Biotechnology | sc-7298 |
| HSC70-HRP conjugated | Santa Cruz Biotechnology | sc-7298-HRP |
| Hydroxy-HIF1a | Cell Signaling | 3434 |
| Lamin B1 | Cell Signaling | 12586 |
| Malic Enzyme 2 | Cell Signaling | 15506S |
| MTHFD2 | Cell Signaling | 98116S |
| Myc | Cell Signaling | 13987S |
| NADK2 | Abcam | ab181028 |
| PHGDH | Cell Signaling | 66350S |
| Phospho-AMPKa (Thr172) | Cell Signaling | 2535 |
| Phospho-c-Myc (Ser62) | Cell Signaling | 13748S |
| Phospho-c-Myc (Thr58) | Cell Signaling | 46650 |
| PYCR1 | Cell Signaling | 37635S |
| PYCR2 | Sigma-Aldrich | HPA056873-100UL |
| SHMT2 | Cell Signaling | 33443S |
| TAZ (D3I6D) Rabbit mAb | Cell Signaling | 70148 |
| TFAM (D5C8) Rabbit mAb | Cell Signaling | 8076 |
| Thymidylate Synthase/TYMS | Cell Signaling | 9045S |
| TUFM | ThermoFisher | PA5-27511 |
| Vimentin | BD biosciences | 550513 |
| YAP (D8H1X) XP® Rabbit mAb | Cell Signaling | 14074 |
| YAP1 (Ser94) | Abbiotec | 254542 |
| Secondary antibody and other reagents | Source | Identifier/catalog number |
| IRDye® 680RD Goat anti-Mouse IgG (H + L) | LICOR | 926-68070 |
| IRDye® 800CW Goat anti-Rabbit IgG (H + L) | LICOR | 926-32211 |
| Goat Anti-Mouse IgG (H+L)-HRP Conjugate | Bio-Rad | 172-1011 |
| Goat Anti-Rabbit IgG (H+L)-HRP Conjugate | Bio-Rad | 172-1019 |
| Bio-Rad colorimetric assay | Bio-Rad | 500-0006 |
| cOmplete Protease Inhibitor Cocktail Tablets | Sigma-Aldrich | 11836153001 |
| Criterion TGX 4-20% 18 well gel | Bio-Rad | 567-1094 |
| Laemmli sample buffer | Bio-Rad | 161-0737 |
| SuperSignal™ West Femto Maximum Sensitivity Substra | ThermoFisher Scientific | 34096 |
| SuperSignal™ West Pico PLUS Chemiluminescent Sub | ThermoFisher Scientific | 34578 |
| Immobilon-FL Polyvinylidene fluoride | Millipore | IPFL00010 |
| Immobilon-P PVDF Membrane | Millipore | IPVH00010 |

### Supplementary Table S1 (3/3)

| Reagents used for cell culture, mammosphere assays | | |
|---|---|---|
| Reagent/Kit | Source | Identifier/catalog number |
| DMEM, high glucose | ThermoFisher Scientific | 11965118 |
| DMEM/F12 medium | ThermoFisher Scientific | 11320082 |
| hydrocortisone | Sigma-Aldrich | H0888 |
| insulin | Sigma-Aldrich | I9278-5ML |
| RPMI 1640 medium | ThermoFisher Scientific | 11875119 |
| RPMI 1640 medium, no L-glutamine | ThermoFisher Scientific | 21870076 |
| sodium pyruvate | Sigma-Aldrich | S8636-100ML |
| uridine | Sigma-Aldrich | U3003-50g |
| B-27 supplement | ThermoFisher | 12587-010 |
| Basic fibroblast growth factor (bFGF) | Sigma-Aldrich | F0291-25uG |
| Corning Ultra low attachment plates 24well | Sigma-Aldrich | CLS3473-24EA |
| Costar® 6-well Clear Flat Bottom Ultra-Low Attachment Multiple Well Plates | Corning | 3471 |
| DMEM/F12 mammosphere assay media | ThermoFisher | 11320082/11320033 |
| SingleQuots | LONZA | CC-4136 |

| Materials for Flow Cytometry | | |
|---|---|---|
| Reagent/Material | Source | Identifier/catalog number |
| ALDEFLUOR™ DEAB Reagent | STEMCELL Technologies, Inc. | 1705 |
| ALDEFLUOR™ Kit | STEMCELL Technologies, Inc. | 1700 |
| LIVE/DEAD™ Fixable Blue Dead Cell Stain Kit | ThermoFisher | L34961 |
| LIVE/DEAD™ Fixable Aqua Dead Cell Stain Kit | ThermoFisher | L34966 |
| Cell Strainer | BD Falcon | 352235 |
| **Seahorse assays** | | |
| Reagent/Material | Source | Identifier/catalog number |
| Seahorse XF Real Time ATP Rate Assay Kit | Agilent | 103592-100 |
| Seahorse XFe24 FluxPak | Agilent | 102340-100 |
| Seahorse XF DMEM | Agilent | 103575-100 |
| Seahorse XF 1.0 M glucose solution | Agilent | 103577-100 |
| Seahorse XF 100 mM pyruvate solution | Agilent | 103578-100 |
| Seahorse XF 200 mM glutamine solution | Agilent | 103579-100 |
| **Tumor dissociation** | | |
| Reagent/Material | Source | Identifier/catalog number |
| Protocol | https://www.miltenyibiotec.com/US-en/applications/all-protocols/Isolation-of-xenografted-cells-from-tumors-by-depletion-of-mouse-cells.html?countryRedirected=1 | |
| Tumor Dissociation Kit, human | Miltenyi Biotec (Bergisch Gladbach, Germany) | #130-095-929 |
| gentleMACS™ Octo Dissociator with Heaters | Miltenyi Biotec | #130-096-427 |
| gentleMACS C Tubes | Miltenyi Biotec | #130-093-237 |
| MACS® SmartStrainers (70 µm) | Miltenyi Biotec | #130-098-462 |
| MACSmix™ Tube Rotator | Miltenyi Biotec | #130-090-753 |
| Mouse Cell Depletion Kit | Miltenyi Biotec | #130-104-694 |
| QuadroMACS™ Starting Kit (LS) | Miltenyi Biotec | #130-091-051 |
| MACS BSA Stock Solution | Miltenyi Biotec | #130-091-376 |
| Pre-Separation Filters (70 µm) | Miltenyi Biotec | #130-095-823 |
| **Others** | | |
| Equipment | Source | |
| Cytation 1 | BioTek | |
| BD FACSVerse Flow Cytometer | BD biosciences (San Jose, CA) | |
| Cellometer K2 | Nexcelom Bioscience (Lawrence, MA) | |
| Seahorse XFe24 Extracellular Flux Analyzer | Agilent Technologies (Santa Clara, CA) | |
| SpectraMax$^{R}$ iD3 microplate reader | Molecular Devices, LLC (San Jose, CA) | |
| Software | Source | URL |
| FlowJo | FlowJo, LLC (Ashland, OR) | |
| Seahorse XF Real-Time ATP Rate assay Report Generator Software | Agilent | https://www.agilent.com/en/products/cell-analysis/xf-real-time-atp-rate-assay-report-generator |
| Extreme Limiting Dilution Analysis (ELDA) | | http://bioinf.wehi.edu.au/software/elda/ |
| Ingenuity Pathway Analysis | Qiagen | https://digitalinsights.qiagen.com/plugins/ingenuity-pathway-analysis/ |
| MetaCore | Clarivate™ | https://portal.genego.com/ |
| Gene Set Enrichment Analysis | UC San Diego & Broad Institute | https://www.gsea-msigdb.org/gsea/index.jsp |
| A custom computational pipeline to determine editing rate (for CRISPR CLPP KO) | | http://github.com/rajchari2/ngs_amplicon_analysis |
